# Motor cortical areas facilitate schema-mediated integration of new motor information into memory

**DOI:** 10.1101/2025.08.05.668621

**Authors:** S. Reverberi, N. Dolfen, B.R. King, G. Albouy

## Abstract

New information is rapidly learned when it is compatible with pre-existing knowledge, i.e. with a previously acquired schematic representation of the learned information. The influence of pre-established schema on learning has been extensively studied in the declarative memory domain, where it was shown that schema-compatible information could be rapidly assimilated into neocortical storage, bypassing the slow hippocampo-neocortical memory transfer process. Schema-mediated learning was recently examined in the motor memory domain; however, its neural substrates remain unknown. The goal of this study was to address this knowledge gap using both univariate and multivariate analyses of functional Magnetic Resonance Imaging (fMRI) data acquired in 60 young healthy participants during the practice of a motor sequence that was either compatible or incompatible with a previously acquired cognitive-motor schema. Consistent with previous literature, our behavioural results suggest that performance of sequential movements was enhanced when practice occurred in a context that was compatible with the previously acquired schema. Brain imaging results show that practice in a schema-compatible context specifically recruited the left primary motor cortex and resulted in a decrease in connectivity between the bilateral motor cortex and a set of task-relevant brain regions including the hippocampus, striatum, and cerebellum. Temporally fine-grained MRI analyses revealed that multivoxel activation patterns in the primary motor and the premotor cortices were modulated by schema-compatibility, with greater pattern similarity detected for sequence elements corresponding to and surrounding novel sequential movements under schema-compatible compared with -incompatible conditions. Altogether, these results suggest that motor cortical regions facilitate schema-mediated integration of novel movements into memory.

## 1. Introduction

The overlap of multiple memory traces related in content is believed to result in the development of a cognitive schema, defined as an associative knowledge structure consisting of the gist representation of previous experiences (Lewis & Durrant, 2011). The availability of an established schema has been demonstrated to facilitate the learning of new information that is compatible with this previous knowledge (van Kesteren et al., 2012). Experimental evidence supporting the schema model of memory consolidation initially came from rodent work (Tse et al., 2007, 2011). After learning a specific pattern of flavour-location associations in an arena, rats were subsequently able to rapidly consolidate novel associations if they were compatible (but not if they were incompatible) with the previously learned set of flavour-location associations (Tse et al., 2007). Schema-facilitated learning was subsequently demonstrated in humans in the declarative memory domain. Memory performance was consistently shown to be enhanced when learning occurred in schema-compatible – compared to incompatible – conditions for a wide range of stimuli including object-scene associations (van Kesteren et al., 2013), face-location and face-house associations (Atienza et al., 2011; Liu et al., 2018), item-color associations (Cycowicz et al., 2008), noun-adjective associations (Bein et al., 2015), auditory-visual associations (Heikkilä et al., 2015), study-related facts (van Kesteren et al., 2014) and movie clips (Keidel et al., 2018; van Kesteren et al., 2010).

Several neurobiological models of schema-mediated learning have been proposed in the declarative memory domain (Lewis & Durrant, 2011; Sekeres et al., 2024; van Kesteren et al., 2012). These models attempt to reconcile the rapid memory consolidation observed under schema-compatible conditions with the standard systems consolidation model suggesting a gradual and slow consolidation process which can span days to years (Squire & Alvarez, 1995). According to this standard model of consolidation, the hippocampus is necessary for the initial acquisition of novel memories, that are then slowly transferred to long-term neocortical storage sites, becoming independent of hippocampal activation (Squire & Alvarez, 1995). Schema models of consolidation propose that this slow transfer process can be bypassed when the novel information to be learned is compatible with previously acquired knowledge (Lewis & Durrant, 2011; Sekeres et al., 2024; van Kesteren et al., 2012). Consistent with this view, neuroimaging studies have demonstrated that the encoding of schema-compatible memories results in increased activity in the medial prefrontal cortex (mPFC) while learning schema-incompatible information recruits the hippocampus (Bonasia et al., 2018; van Kesteren et al., 2013). In the same vein, both the mPFC (Audrain & McAndrews, 2022) and the angular gyrus (Wagner et al., 2015) have been shown to play critical roles in the retrieval of schema-compatible memories and rule-based schema memories. Last, in line with the idea that the integration of novel memories compatible with pre-existing schema bypasses slow hippocampo-cortical transfer of information, connectivity studies have shown that the integration of schema-compatible, versus incompatible, information resulted in reduced functional connectivity between the hippocampus and the mPFC (van Kesteren et al., 2010, 2014).

The schema model of memory consolidation was only more recently examined in the non-declarative memory domain. For example, a study on perceptual memory showed that the learning of new melodies was enhanced for participants of Western culture when the melodies conformed to a tone distribution pattern commonly found in Western music (Durrant et al., 2015). In the motor memory domain, it has been proposed that schematic representations of motor sequences can develop after learning, and that the corresponding cognitive-motor schema encompasses the association between the performed movements and their ordinal position within the sequence stream (King et al., 2019). In line with observations from other memory domains, previous work demonstrated that a novel sequence whose ordinal structure was compatible with the previously acquired motor sequence schema was learned significantly faster than a schema-incompatible sequence (King et al., 2019). Overall, evidence from the behavioral studies reviewed above suggests that the schema model of memory presents similar characteristics across memory domains. However, it remains unknown whether this similarity extends to the underlying neural processes, as the cerebral substrates subtending schema-mediated learning have only been investigated in the declarative memory domain.

The goal of the current investigation was therefore to examine the neural processes supporting schema-mediated learning in the motor memory domain. To do so, we used both univariate and multivariate analyses of functional Magnetic Resonance Imaging (fMRI) data acquired during the learning of novel motor sequences that were either compatible or incompatible with a pre-existing cognitive-motor schema. Our overarching hypothesis was that the learning of schema-incompatible motor sequences would recruit brain areas traditionally involved in the *learning of new motor sequences* and would therefore rely on increased activity and connectivity in the cerebellum, striatum and the hippocampus (as well as their cortical projections; Albouy et al., 2013; Dayan & Cohen, 2011; Doyon et al., 2003). In contrast, we expected that learning schema-compatible sequences would recruit regions involved in *the cortical storage of previously learned motor sequences* such as the primary motor cortex (M1, Penhune & Steele, 2012) as well as the premotor and parietal cortices (Berlot et al., 2020; Yokoi & Diedrichsen, 2019) and would be accompanied by decreased connectivity between these storage cortical regions and the brain areas involved in novel motor learning described above.

## 2. Methods

### 2.1. Participants

Sixty healthy young volunteers (mean age: 23; age range: 19-30 years old) were recruited from KU Leuven and surroundings. The following criteria were used to determine participant inclusion: 1) right-handed, as assessed with the Edinburgh Handedness Inventory (Oldfield, 1971), 2) no prior extensive training with a musical instrument requiring dexterous finger movements (e.g., piano, guitar) or as a professional typist, 3) free of medical, neurological, psychological, or psychiatric conditions, including depression and anxiety as assessed by the Beck’s Depression and Anxiety Inventories (Beck et al., 1988, 1996), 4) no indications of abnormal sleep, as assessed by the Pittsburgh Sleep Quality Index (PSQI (Buysse et al., 1989)), 5) not considered extreme morning or evening types, as quantified with the Horne & Ostberg chronotype questionnaire (Horne & Ostberg, 1976), 6) free of psychoactive and sleep-influencing medications, 7) non-smokers, and 8) not having completed trans-meridian trips or worked night shifts in the month prior to participation. All participants gave written informed consent before the start of the study and were compensated for their participation. Participants were pseudo-randomly assigned to the schema-compatible (COMP) or schema-incompatible (INCOMP) group, with an equal gender ratio across the groups (see supplementary Table S1 for participants demographics).

#### 2.1.1. Sample size justification

Our sample size was based on a main effect of group (COMP vs. INCOMP) measured on the response time for novel transitions in our previous behavioural experiment employing a similar experimental design (King et al., 2019). This group main effect (F(1,36)=5.29, *p*=0.027, Ƞp^2^=0.128, Cohen’s F=0.383) resulted in an estimated minimum inclusion of 21 subjects per group, as assessed via G*Power (Faul et al., 2007) (Effect size f=0.383, tails=2; alpha=0.05, power=0.80, correlation among repeated measures=0.72). As we expected performance measured in the MRI scanner to be more variable than when measured outside the scanner in a behavioural study, we aimed for an increased sample size of 30 subjects per group (60 participants in total).

### 2.2. Motor task

Participants performed an explicit serial reaction time task (SRTT; Nissen & Bullemer, 1987) coded and implemented in MATLAB (Mathworks Inc., Sherbom, MA) using Psychophysics Toolbox version 3 (Kleiner et al., 2007). Participants performed the task either on a computer keyboard while seated in front of a laptop screen (Session 1, see experimental procedure) or lying on their back in the MRI scanner, on a specialized MR-compatible keyboard (Session 2). In both sessions, the participants were not able to see their fingers.

During the task, participants were presented with 8 squares on the computer screen, corresponding spatially to the 8 fingers used to perform the task (thumbs were not used, see Figure 1). The outline of the squares was coloured red or green during periods of rest or practice, respectively. During practice, cues (full green squares, see Figure 1) consecutively appeared in the different spatial locations on the screen, and participants were instructed to press the corresponding key as fast and as accurately as possible. The order of these visual cues followed either a pseudorandom pattern (random SRTT) or a deterministic 8-element sequence pattern (sequential SRTT). In the latter case, participants were informed that the order of key presses would follow a sequential pattern but were not given any additional information about the sequence (e.g., sequence length, number of repetitions). For the pseudorandom condition, each of the eight keys/fingers was used once every 8 keypresses, ensuring that the distribution of keys/fingers was the same as in the sequence condition.

The number of key presses per practice block (64 vs. 96), the duration of the rest blocks (15s vs. 10s) and the response-stimulus interval (RSI=0s vs. 2s [jittered between 1.5-2.5s]) were different when the task was performed inside or outside the scanner (see *Experimental design*) in order to optimize the analyses of the MRI data (see *Experimental design* and *Multivariate analyses* sections below).

### 2.3. Experimental design

The experimental design was similar to our previous work (King et al., 2019; Reverberi et al., 2023) and is presented in Figure 1. Participants completed two sessions of the serial reaction time task separated by a delay of approximately 24 hours. All participants were instructed to follow a regular sleep/wake schedule according to their own preferred rhythm (i.e., ±1h of their habitual bed and wake time with bedtime no later than 1AM and a minimum 7h of sleep per night) for the 3 nights prior to Session 1 and the night between the 2 experimental sessions. Compliance to this schedule was assessed with sleep diaries for the 3 nights leading up to the experiment and with wrist actigraphy recordings (ActiGraph, Pensacola, USA) for the night between the 2 experimental sessions. Sleep quantity and quality for the nights preceding each experimental session were also assessed via the St. Mary’s sleep questionnaire (Ellis et al., 1981). Participants were instructed to avoid alcohol, caffeine, and intensive exercise during the 12h preceding each experimental session. At the start of each session, vigilance was estimated with the psychomotor vigilance task (PVT, Dinges & Powell, 1985). Results related to sleep and vigilance data are reported in supplementary Table S1.

During Session 1 (S1), that took place outside the scanner, all participants learned the same motor sequence (Sequence 1; 4-7-3-8-6-2-5-1, in which 1 and 8 represent the left and right little fingers, without the thumbs). During Session 2 (S2), in line with our previous research (King et al., 2019; Reverberi et al., 2023), all participants learned a new motor sequence that was presented in an ordinal context that was either highly compatible (COMP group) or highly incompatible (INCOMP group) with the cognitive-motor schema established during Session 1 and consisting of the ordinal structure of the previously learned Sequence 1 (King et al., 2019). Specifically, the sequence performed during Session 2 (S2) was identical to Sequence 1 with the exception that the position of 2 keys was switched (keys 2 and 3, “novel” keys in-text and highlighted in red in Figure 1). The starting point of the S2 sequences was different between groups such that the sequences were performed either in a highly compatible (COMP) or incompatible (INCOMP) ordinal framework. Specifically, the ordinal structure of the S2 sequence (4-7-2-8-6-3-5-1) in the COMP group was highly compatible to that of Sequence 1 as 75% of the keys were presented in the same ordinal position as S1 and thus 75% of key/ordinal position pairings learned in Sequence 1 were preserved. In contrast, in the INCOMP group, the S2 sequence (8-6-3-5-1-4-7-2) contained only 12.5% of keys that were presented in the same ordinal position as S1 (i.e. a single key/ordinal position pairing was preserved from Sequence 1).

In each session, the sequence task was divided into multiple sections. During S1, participants first performed 4 blocks of a pseudo-random SRTT (*random*) to assess general motor execution. This was followed by the execution of a sequential SRTT (*training*: 20 blocks, *slow*: 2 blocks, and *test*: 4 blocks). In the training and test blocks, the task was performed with RSI=0s with a design that is identical to that of our previous work (King et al., 2019; Reverberi et al., 2023). In the slow blocks, RSI was set to ∼2s (jittered between 1.5-2.5s) to familiarize participants with the task design used in the MRI scanner the subsequent day (see below). The test blocks were administered approximately 1 minute after the slow runs to assess end-of-training performance following the dissipation of mental and physical fatigue (Pan & Rickard, 2015). During S2, participants performed 3 sections of the sequential SRTT. These included a *pre-test* outside the scanner (4 blocks), which was followed by 8 runs of task performed in the MRI scanner (*MRI*), and finally a *post-test* including 8 practice blocks outside the scanner. Pre- and post-tests were performed with RSI=0s to assess performance outside the scanner with a similar design as during S1, i.e., without the timing constraints of the MRI session. The task in the scanner was performed with a slower RSI of ∼2s (jittered between 1.5-2.5s) to optimize event-related MRI analyses.

For the sections of task referred to as *random, training, test, pre-test* and *post-test* (see Figure 1), performed outside of the MRI scanner in Sessions 1 and 2, each block of practice contained 64 key presses (corresponding to 8 repetitions of the 8-element sequence or eight pseudo-random key presses) and was separated from the next block by a rest period of 15 seconds. The sections of task referred to as *slow* in Session 1 and *MRI* in Session 2 contained 2 and 8 practice runs, respectively, with each run including 96 keypresses (corresponding to 12 repetitions of the sequence) and three 10-second rest periods randomly inserted between sequence repetitions to minimize fatigue.

**Figure 1:**
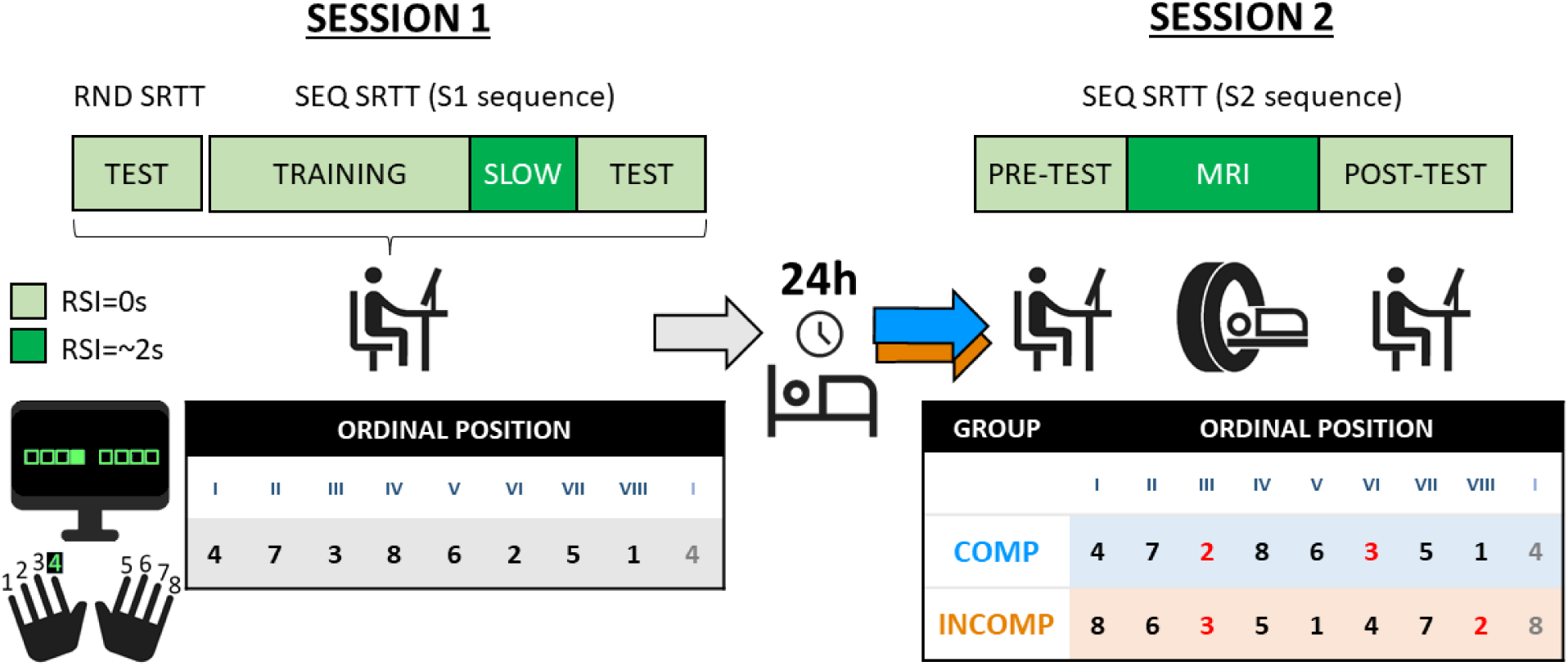
Task and procedure. Participants completed two experimental sessions separated by 24 hours, in which they performed different SRT tasks. In Session 1 (S1), participants were seated at a desk and completed a pseudo-random SRTT (RND) test to assess baseline performance, followed by the sequential SRTT (SEQ; divided into training, slow, and test sections), during which the cognitive-motor schema of S1 sequence was acquired. Participants were instructed to press the key shown on-screen with the corresponding finger as fast and as accurately as possible. Participants learned the sequence 4-7-3-8-6-2-5-1 (1 to 8: left to right little finger) shown in the table, with the roman numerals (I to VIII) representing the ordinal position of the keys/fingers to be pressed in the sequence. In Session 2, participants learned a new motor sequence (referred to as S2 sequence). They performed this sequential SRTT during a pre-test, followed by the MRI session, and then a post-test (pre- and post-tests were performed outside the MRI scanner). The S2 sequence was created by switching two keys of the S1 sequence (referred to as “novel” keys, shown in red in S2 table). The compatibility to the previously learned ordinal structure was manipulated by changing the starting point of the S2 sequence between the different groups such that the sequence was performed in an either highly compatible or incompatible ordinal framework in the COMP and INCOMP groups, respectively (see section “Experimental design” for details). SRT tasks were performed with a response to stimulus interval (RSI) of 0s (light green), or of 2s on average (dark green); the latter being used to optimize the MRI analyses (see section “FMRI multivariate analyses” for details). Icons used in this figure were adapted from Google Material Symbols (Apache License, version 2.0).

### 2.4. FMRI acquisition and preprocessing

FMRI images were acquired using a Phillips Achieva 3.0T MRI System equipped with a 32-channel head coil. Structural T1-weighted images were acquired with a 3D MP-RAGE sequence (TR=9.5ms, TE=4.6ms, TI=858.1ms, FA=9°, 160 slices, FoV=250×250mm^2^, matrix size=256×256×160, voxel size=0.98×0.98×1.20mm^3^). Functional images were acquired during the 8 task runs with a T2* gradient echo-planar sequence using axial slice orientation that covers the whole brain (TR=2000ms, TE=30 ms, FA=90°, 54 transverse slices, 2.5mm slice thickness, 0.2 mm inter-slice gap, FoV=210×210mm^2^, matrix size=84×82×54 slices, voxel size=2.5×2.56×2.5mm^3^).

Functional data were preprocessed using Statistical parametric mapping (SPM12; Welcome Department of Imaging Neuroscience, London, UK) implemented in MATLAB. Functional images were first slice-time corrected using the middle slice as reference slice. Functional time series were then realigned with a two-step approach using rigid body transformations iteratively optimized to minimize the residual sum of squares between each functional image and the first image of its corresponding run in a first step, and between each functional image and the across-run mean functional image in a second step. Analysis of the realignment parameters across all included runs indicate minimal head movement during scanning, as the mean maximum translation in the three directions in our sample was of 0.46±0.28mm in the COMP group and of 0.41±0.30mm in the INCOMP (no difference between groups, independent t-test *t*=0.54, *p*=0.59). Note that imaging runs were excluded from data analyses if the maximum movement within the run exceeded ∼2 voxels (i.e., 5mm) in any of the three dimensions (see section Data exclusions for details of excluded runs). Preprocessing further included co-registration of the pre-processed functional images to the structural T1-image using rigid body transformation optimized to maximize the normalized mutual information between the functional and anatomical images. The T1 image was then segmented into grey matter, white matter, cerebrospinal fluid, bone, soft tissue, background, and each participant’s forward deformation field was used for the normalization step (univariate analyses only). For the univariate analyses, structural and functional data were then normalized to MNI space (resampling size of 2x2x2mm) and functional data were spatially smoothed (isotropic Gaussian kernel, 8 mm full-width half-maximum). For the multivariate analyses, data was maintained in native space to account for inter-individual variability in the topography of memory representations (Haxby et al., 2001).

### 2.5. Statistical analyses

#### 2.5.1. Behavioural analyses

All statistical analyses on behavioural data were performed using IBM SPSS Statistics for Windows, version 28 (IBM Corp, 2021). In case of violation of the sphericity assumption, we applied Greenhouse-Geisser corrections for ε≤0.75, Huynh-Feldt corrections for ε>0.75 (J. P. Verma, 2015).

##### 2.5.1.1. Behavioural data exclusions

Due to experimental computer malfunction resulting in missing task blocks, (i) one participant of the INCOMP group was excluded from the analyses of the Session 1 training data (Session 1 training analyses therefore included 59 participants, 30 in the COMP and 29 in the INCOMP group); (ii) another participant of the INCOMP group was excluded from the analyses of the Session 1 test data (which therefore included 59 participants, 30 in the COMP and 29 in the INCOMP group); and (iii) one participant of the COMP group was excluded from the analyses of the Session 2 pre-test data (which therefore included 59 participants, i.e., 29 in the COMP and 30 in the INCOMP group). One participant in the INCOMP group was excluded from the analyses of the behavioural data collected in the MRI scanner as they did not follow task instructions during one run of the task (large lags between responses due to drowsiness; Session 2 MRI behavioural analyses therefore included 59 participants, 30 in the COMP and 29 in the INCOMP group). Note that this run was also excluded from the MRI analyses (see *MRI data exclusions*).

##### 2.5.1.2. Primary outcome variables

The primary outcome variables for the SRTT analyses were performance speed measured as the response time (RT), i.e. the time between cue presentation and participant key press, and performance accuracy, i.e. the percentage of correct key presses. The averaged response times for correct keypresses and the accuracy were computed for each block or run of practice on the task. Performance on the sequential SRTT was analysed with repeated measures ANOVAs separately for each section of the task (see *Motor task* for details) using within-subject factor *block* or *run,* between-subject factor *group* (COMP/INCOMP), and dependent measure *RT* or *accuracy*.

##### 2.5.1.3. Negative control analyses

Negative control analyses were performed on vigilance scores (i.e., mean reaction time on the PVT task), sleep quantity (hours) and quality (score 1-5), inclusion questionnaire scores (see below for details), and pseudo-random SRTT data (speed and accuracy measures). Group differences in sleep quantity and quality, vigilance, as well as handedness, depression, anxiety, and chronotype scores (see section Participants and Table S1 for details) were assessed with independent-samples t-tests (two-sided, equal variances assumed where Levene’s test non-significant). Group differences in the Session 1 pseudo-random SRTT, reflecting general motor execution, were assessed with a *group* x *block* ANOVA (see supplementary Table S2).

#### 2.5.2. FMRI analyses

Univariate statistical analyses were performed with SPM12 (Welcome Department of Imaging Neuroscience, London, UK) implemented in MATLAB. Multivariate pattern analyses were performed using the CoSMoMVPA toolbox (Oosterhof et al., 2016).

##### 2.5.2.1. MRI data exclusions

One participant was excluded from all MRI data analyses due to excessive head motion in over half of the MRI runs, resulting in a final sample size of 59 participants for all MRI analyses with 30 participants for the INCOMP group and 29 for the COMP group. One single run of MR task was additionally excluded from the neuroimaging analyses for two additional participants of the INCOMP group, one due to excessive head motion and one due to non-compliance with task instructions (detailed under *Behavioural data exclusions*) in these particular runs. All other runs were included in the analyses.

##### 2.5.2.2. Univariate analyses

###### 2.5.2.2.1. Activation-based analyses

The univariate analysis of fMRI data collected during Session 2 was performed in two steps. In the first-level fixed effects analyses, one regressor of interest represented task practice and included one event per each key/finger cue that was modelled using stick functions (0ms duration) locked to cue onset and convolved with the canonical hemodynamic response function. Regressors of no interest were also modelled, in particular regressors representing incorrect key presses and key presses performed during rest periods, as well as motion regressors (3 translations and 3 rotations) derived from functional volumes realignment. Finally, the data was high-pass filtered with a cut-off period of 128s to remove low-frequency drifts, and serial correlations in the signal were estimated using an autoregressive (order 1) plus white noise model and a restricted maximum likelihood (ReML) algorithm. Following first-level analyses, the resulting statistical parametric maps representing the contrast of task practice vs. rest were spatially smoothed for each participant (Gaussian kernel, 6mm FWHM) and entered into second-level, random effects analyses. The second-level analyses examined brain responses during task practice versus rest across the two experimental groups using a one-sample t-test (see Supplementary Table S5) as well as the difference in neural activation between the two groups of participants (COMP and INCOMP) using a two-sample t-test (see Results in the main text, and Supplementary Table S6 for activations outside regions of interest).

###### 2.5.2.2.2. Connectivity-based analyses

Task-related functional connectivity was examined using psychophysiological interaction (PPI) analyses. We assessed connectivity of two homologous seed regions showing a significant difference in activation between the COMP and INCOMP groups in the activation-based fMRI analysis described above (i.e., left M1, -34 -32 58mm and right M1, 40 -20 52mm). For each participant, singular value decomposition of the time series was used to extract the first eigenvariate of the signal across the voxels included in a 6mm-radius sphere centered on the peaks of the activation clusters reported at the group level in each contrast of interest (COMP>INCOMP and INCOMP>COMP). General linear models (GLMs) were generated at the individual level and consisted of three regressors representing: (i) the neural activity in the reference area (“physiological” regressor), (ii) the practice of the learned sequence (“psychological” regressor), and (iii) the interaction of interest between the previous two regressors. To build this interaction regressor, the underlying neuronal activity was first estimated by a parametric empirical Bayes formulation, then combined with the psychological regressor and finally convolved with the standard hemodynamic response function (Gitelman et al., 2003). Movement parameters were also included as regressors of no interest. A significant PPI reflects a change in the regression coefficients, thus a change in the strength of the functional interaction, between any reported brain region and the reference seed region, related to the practice of the learned sequence as compared to rest. Similar to the activation-based analyses described above, individual summary statistic images obtained at the first-level (fixed effects) PPI analysis were spatially smoothed (6mm FWHM Gaussian kernel) and entered in a second-level (random-effects) analysis using a two-sample t-test to compare the strength of the functional interaction between the two groups of participants.

###### 2.5.2.2.3. Statistics

The set of voxel values resulting from each second level analysis described above (activation and functional connectivity) constituted maps of the t statistic for each contrast of interest [SPM(T)] and were thresholded for display at *p*<0.001 (uncorrected for multiple comparisons). Statistical inferences were performed at a threshold of *p*<0.05 after family-wise error (FWE) correction for multiple comparisons over small spherical 10mm-radius volumes centered on coordinates derived from previous literature and located within regions of interest (small volume correction of the family-wise error rate (Poldrack, 2007), coordinates used for correction reported in Table S3). Further correction for multiple volumes of interest was performed within each contrast using the Benjamini-Hochberg correction of the false discovery rate (FDR, *p*<0.05) (Benjamini & Hochberg, 1995). Beta values extracted from activation-and connectivity-based analyses were graphically visualized using the MATLAB package *DataViz* (Povilas Karvelis & oyvindlr, 2024).

##### 2.5.2.3. Multivariate analyses

The univariate analyses of the fMRI data described above allowed us to investigate group differences in brain activity/connectivity during task practice as compared to rest. We additionally performed multivariate pattern analyses to provide a more fine-grained analysis of brain responses associated to each sequential movement. This allowed us to investigate the specific effects of schema compatibility on brain responses associated to specific elements of the motor sequence.

###### 2.5.2.3.1. ROI selection

The multivariate analyses followed a procedure analogous to that of Dolfen and colleagues (Dolfen et al., 2024) and used a region of interest (ROI) approach. The following ROIs were selected a priori, given their established involvement in motor sequence learning (Albouy et al., 2013; Berlot et al., 2020; Dayan & Cohen, 2011; Doyon et al., 2003, 2009): the hippocampus, putamen, primary motor cortex (M1), premotor cortex, and anterior superior parietal lobule (aSPL). Additionally, two ROIs derived from the declarative memory schema literature were added for exploratory analyses (i.e., the medial prefrontal cortex and the angular gyrus, results presented in the Supplementary material, section “*Exploratory analyses on declarative memory schema ROIs*”, Table S7, Figure S1). Based on the results of the univariate analyses, the M1 ROI was divided into left and right M1 while all other ROIs were bilateral. All ROIs were anatomically defined. Subcortical ROIs (hippocampus and putamen) were defined in native space using FSL’s automated subcortical segmentation protocol (FIRST, Patenaude et al., 2011). Cortical ROIs were created in MNI space and then converted into native space. The ROIs of M1, premotor cortex, medial prefrontal cortex, and angular gyrus were created using the Brainnettome atlas (Fan et al., 2016). M1 included the upper limb and hand function regions of Brodmann area (BA) 4. The premotor cortex (PMC) was defined as the dorsal (A6cdl; dorsal PMC) and ventral (A6cvl; ventral PMC) part of BA 6. The medial prefrontal cortex mask included the medial regions of BA 10 and BA 14 (A10m, A14m). The angular gyrus included the caudal, rostrodorsal, and rostroventral areas of BA 39 (A39c, A39rd, A39rv). The aSPL ROI was created the Julich brain atlas (Amunts et al., 2020) and was defined to include the anterior, medial, and ventral intraparietal sulcus. Only voxels inside the brain and with at least 10% probability of belonging to grey matter were included in the ROI masks. The average number of voxels within each ROI is reported in Supplementary Table S4.

###### 2.5.2.3.2. Representational similarity analyses

Representational similarity analyses (RSA) were designed to examine the effect of schema compatibility on the coding of information about finger movements in their temporal position in the new sequences. For each ROI, and for each participant, representational similarity values were therefore calculated for each of the 8 sequence elements. To do this, a GLM was created with one regressor per each key/ordinal position pairing. This resulted in a total of 8 regressors of interest per run of task. For each regressor, events of interest were modelled with delta functions (0ms duration) time-locked to cue onset. Keypresses during rest and movement parameters derived from functional volume realignment were considered regressors of no interest. High-pass filtering and serial correlation estimation were performed as for the univariate analyses. The resulting statistical maps allowed the extraction of one vector of t-values (size: number of voxels per ROI) for each sequence element (i.e., each key/ordinal position pairing), each run of task, and participant. T-values were normalized by subtracting the mean t-value across all 8 regressors from each voxel in each ROI, separately for each task run.

Representational similarity analyses were performed following the procedure of Dolfen et al. (2024). Specifically, Pearson correlation coefficients were computed for each regressor by correlating vectors of t-values between runs. To do so, the full dataset of 8 runs was randomly divided into two independent subsets of 4 runs, and t-values were averaged across each subset. This resulted in 2 sets of 8 averaged t-value vectors. Next, correlations were computed between runs for each sequence element (e.g., correlating the t-value vector corresponding to key 4 / ordinal position 1 in the first set of runs and in the second set of runs). This procedure was repeated 70 times (i.e., the number of possible combinations of 8 runs divided in 2 groups of 4) and correlations were then averaged across the 70 iterations, resulting in an averaged similarity value per sequence element, ROI, and participant. Finally, the resulting correlation values were Fisher-transformed. As multivariate patterns are believed to reflect representational content (Mur et al., 2009), the computed similarity values represent the consistency in a brain region’s representation of each sequential element, from low similarity (low consistency of responses across voxels in the brain region, suggesting minimal representation of the stimulus information in the brain region) to high similarity (high consistency of responses across voxels in the brain region, suggesting that the region carries information about this sequential element).

To investigate the effect of schema compatibility on the neural representation of the sequence of movements, the computed similarity values for each ROI were entered in a 2x8 ANOVA with between-subject factor *group* (COMP/INCOMP) and within-subject factor *ordinal position* (1 through 8). The Benjamini-Hochberg false discovery rate (FDR, *p*<0.05) correction (Benjamini & Hochberg, 1995) was used to correct for testing across multiple ROIs (i.e., the 6 ROIs presented in the main text). When a significant interaction of group and ordinal position was detected, follow-up independent samples t-tests (two-sided, equal variances assumed where Levene’s test non-significant) were performed to examine differences in representation for each sequence element between groups. Benjamini-Hochberg FDR correction was then applied over the 8 sequence elements. All statistical analyses on the representational similarity data were performed using IBM SPSS Statistics for Windows, version 28 (IBM Corp, 2021). In case of violation of the sphericity assumption, we applied Greenhouse-Geisser corrections for ε≤0.75, Huynh-Feldt corrections for ε>0.75 (J. P. Verma, 2015). Results of the RSA analyses were graphically visualized using the MATLAB package *DataViz* (Povilas Karvelis & oyvindlr, 2024).

## 3. Results

### 3.1. Behavioural results

The behavioural analyses focused on performance speed (response time, RT) for correct keypresses and on accuracy (% correct responses), averaged across each block or run of practice during the sequential SRTT (Figure 2). See also supplementary Table S2 for similar analyses of the random SRTT showing that general motor execution did not differ between groups at baseline (Figure 2, Random).

<u>Session 1</u>: Performance on the sequential SRT task during S1 is depicted in Figure 2A and the output of the corresponding statistical analyses is presented in Table 1. As expected, participants successfully learned Sequence 1, as evidenced by the significant block effect observed on performance speed across all task sections in S1. Accuracy remained overall high and stable throughout S1. Crucially, no significant group effects or block x group interactions were detected during S1 on either RT or accuracy, indicating that performance did not differ between the two experimental groups prior to the experimental manipulation introduced in S2.

<u>Session 2</u>: Results suggest that participants successfully learned Sequence 2, with a significant effect of block on RT for all task sections, indicating progressively faster task performance (Figure 2B, Table 2). In line with our hypothesized effect of schema compatibility on learning, a significant group difference (F(1,57)=4.99, *p*=0.03) was detected on response time during the pre-test administered prior to and outside the MR scanner, with the COMP group performing significantly faster than the INCOMP group. These results are in line with our previous research (King et al., 2019) and show an overall performance advantage when practice takes place in an ordinal framework that is compatible – as compared to incompatible – with the previously learned sequence.

This group difference, however, did not persist during the 8 runs of practice performed in the MR scanner, when task practice was slow-paced to accommodate imaging requirements, nor during the post-test performed outside of the scanner (i.e., in the same conditions as the pre-test). No group differences or block x group interactions were observed on the accuracy measure.

**Figure 2:**
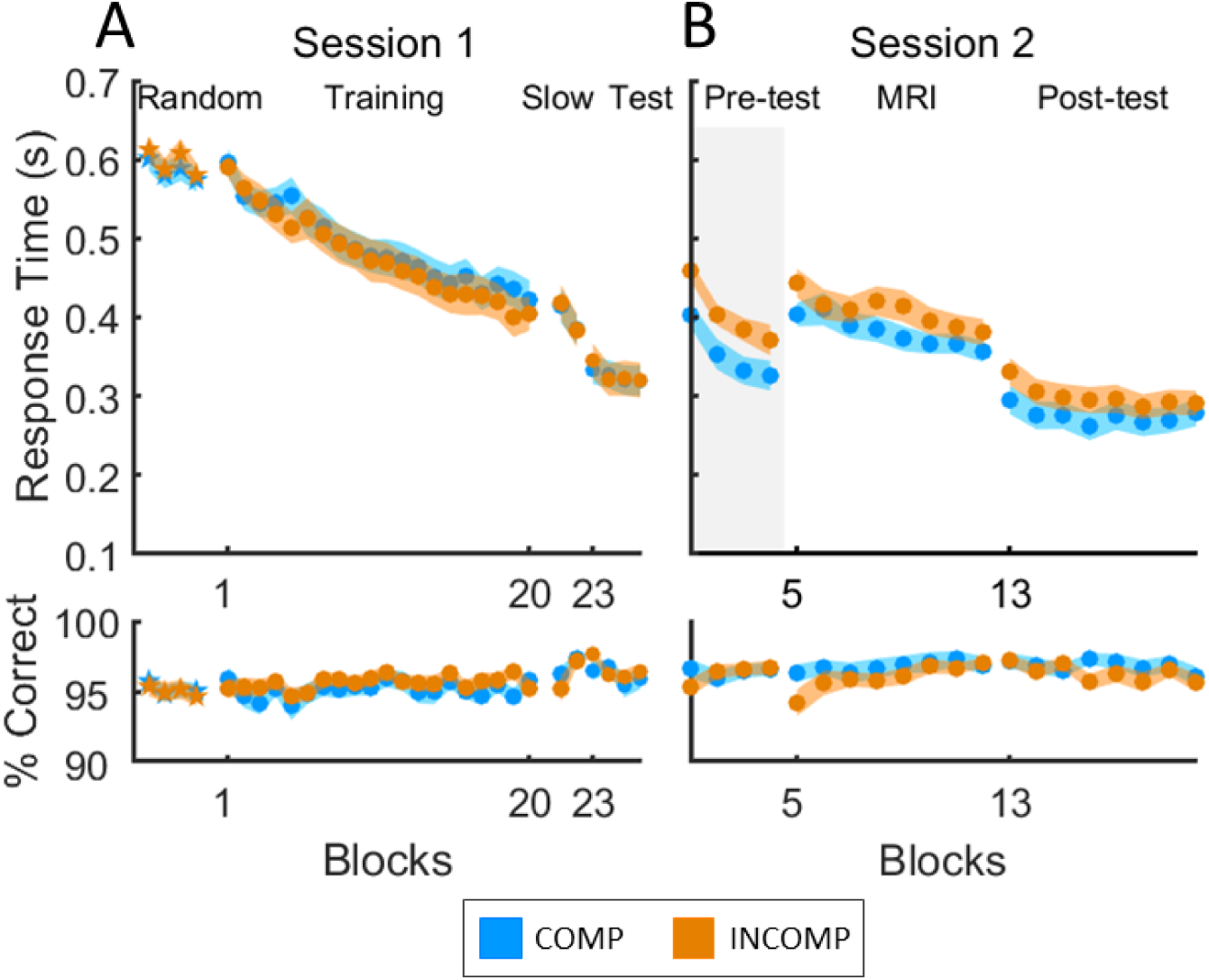
Mean response time (RT, in seconds, top panels) and accuracy (% correct keys, bottom panels) per block are shown separately for the COMP group (blue; N=29 for S2 Pre-test, N=30 for all other sections) and INCOMP group (orange; N=29 for S1 Training, S1 Test, and S2 MRI, N=30 for all other sections) during Session 1 (A) and Session 2 (B). Session 1 included both random (star markers) and sequential (circle markers) SRTT practice. Session 1 started with 4 blocks of random SRTT (response to stimulus interval, RSI=0s) and was followed by 20 blocks of training (RSI=0s), 2 blocks of slow practice (RSI=∼2s) and 4 blocks of test (RSI=0s) on the sequential SRTT. Session 2 consisted of 8 blocks of practice performed in the MR scanner (RSI=∼2s) preceded by 4 blocks of pre-test (RSI=0s) and followed by 8 blocks of post-test (RSI=0s) performed outside the scanner. Shaded coloured areas represent the standard error of the mean. A shaded grey rectangle highlights the section of the task for which a significant difference in performance was detected between groups.

**Table 1:** Sequential SRT task performance during Session 1.

| RT |  |  |  |  | Accuracy |  |  |  |
| --- | --- | --- | --- | --- | --- | --- | --- | --- |
| Effect | df | F | p | Part $\eta^2$ | df | F | p | Part $\eta^2$ |
| *Session 1 Training* |  |  |  |  |  |  |  |  |
| Block | 7.63,434.72 | 67.91 | **<0.001\*** | .544 | 12.88,734.37 | 1.28 | 0.22 | 0.022 |
| Group | 1,57 | 0.15 | 0.70 | 0.003 | 1,57 | 0.76 | 0.39 | 0.013 |
| B x G | 7.63,434.72 | 0.96 | 0.46 | 0.017 | 12.88,734.37 | 0.54 | 0.90 | 0.009 |
| *Session 1 Slow* |  |  |  |  |  |  |  |  |
| Block | 1,58 | 13.79 | **<0.001\*** | 0.192 | 1,58 | 10.24 | **0.002\*** | 0.150 |
| Group | 1,58 | 0.00 | 0.96 | 0.000 | 1,58 | 0.72 | 0.40 | 0.012 |
| B x G | 1,58 | 0.07 | 0.79 | 0.001 | 1,58 | 0.85 | 0.36 | 0.014 |
| *Session 1 Test* |  |  |  |  |  |  |  |  |
| Block | 3,171 | 4.30 | **0.01\*** | 0.070 | 3,171 | 2.98 | **0.03\*** | 0.050 |
| Group | 1,57 | 0.01 | 0.94 | 0.000 | 1,57 | 0.45 | 0.51 | 0.008 |
| B x G | 3,171 | 0.81 | 0.49 | 0.014 | 3,171 | 1.29 | 0.28 | 0.022 |

Significant values are marked with an asterisk. RT = response time; df = degrees of freedom; Part $\eta^2$ = partial eta squared; BxG = block x group interaction.

**Table 2:** Sequential SRT task performance during Session 2.

| RT |  |  |  |  | Accuracy |  |  |  |
| --- | --- | --- | --- | --- | --- | --- | --- | --- |
| Effect | df | F | p | Part $\eta^2$ | df | F | p | Part $\eta^2$ |
| Session 2 Pre-test |  |  |  |  |  |  |  |  |
| Block | 2.15,122.32 | 93.86 | <0.001* | 0.622 | 3,171 | 1.03 | 0.38 | 0.018 |
| Group | 1,57 | 4.99 | 0.03* | 0.080 | 1,57 | 0.06 | 0.80 | 0.001 |
| B x G | 2.15,122.32 | 0.38 | 0.70 | 0.007 | 3,171 | 1.76 | 0.16 | 0.030 |
| Session 2 MRI |  |  |  |  |  |  |  |  |
| Block | 4.39,250.12 | 12.18 | <0.001* | 0.176 | 4.18,238.24 | 3.68 | 0.01* | 0.061 |
| Group | 1,57 | 1.55 | 0.22 | 0.026 | 1,57 | 1.17 | 0.29 | 0.020 |
| B x G | 4.39,250.12 | 1.21 | 0.31 | 0.021 | 4.18,238.24 | 1.23 | 0.30 | 0.021 |
| Session 2 Post-test |  |  |  |  |  |  |  |  |
| Block | 4.19,242.98 | 8.23 | <0.001* | 0.124 | 6.47,375.23 | 1.60 | 0.14 | 0.027 |
| Group | 1,58 | 1.17 | 0.28 | 0.020 | 1,58 | 0.95 | 0.33 | 0.016 |
| B x G | 4.19,242.98 | 0.98 | 0.42 | 0.017 | 6.47,375.23 | 1.03 | 0.41 | 0.017 |

Response Time (RT) and Accuracy measures are provided for all keys (A), learned keys (B), and novel keys (C). Df = degrees of freedom; Part $\eta^2$ = partial eta squared; BxG = block x group interaction. Significant p-values are marked with an asterisk and bold font.

### 3.2. Neuroimaging results

#### 3.2.1 Univariate analyses

To assess the effect of schema compatibility on overall task-related brain activity, we compared the practice versus rest contrast between the COMP and INCOMP groups (and see supplemental Table S5 for task-related brain activation across groups). Results show that activity in the left M1, left SMA, left precuneus, right hippocampus, and right cerebellum was significantly greater during performance of a schema-compatible, compared to incompatible, sequence (COMP > INCOMP, Figure 3, corresponding results presented in Table 3A). In contrast, activity in the right M1 was greater in a schema-incompatible, compared to compatible, context (INCOMP > COMP, Figure 3, Table 3B).

**Figure 3:**
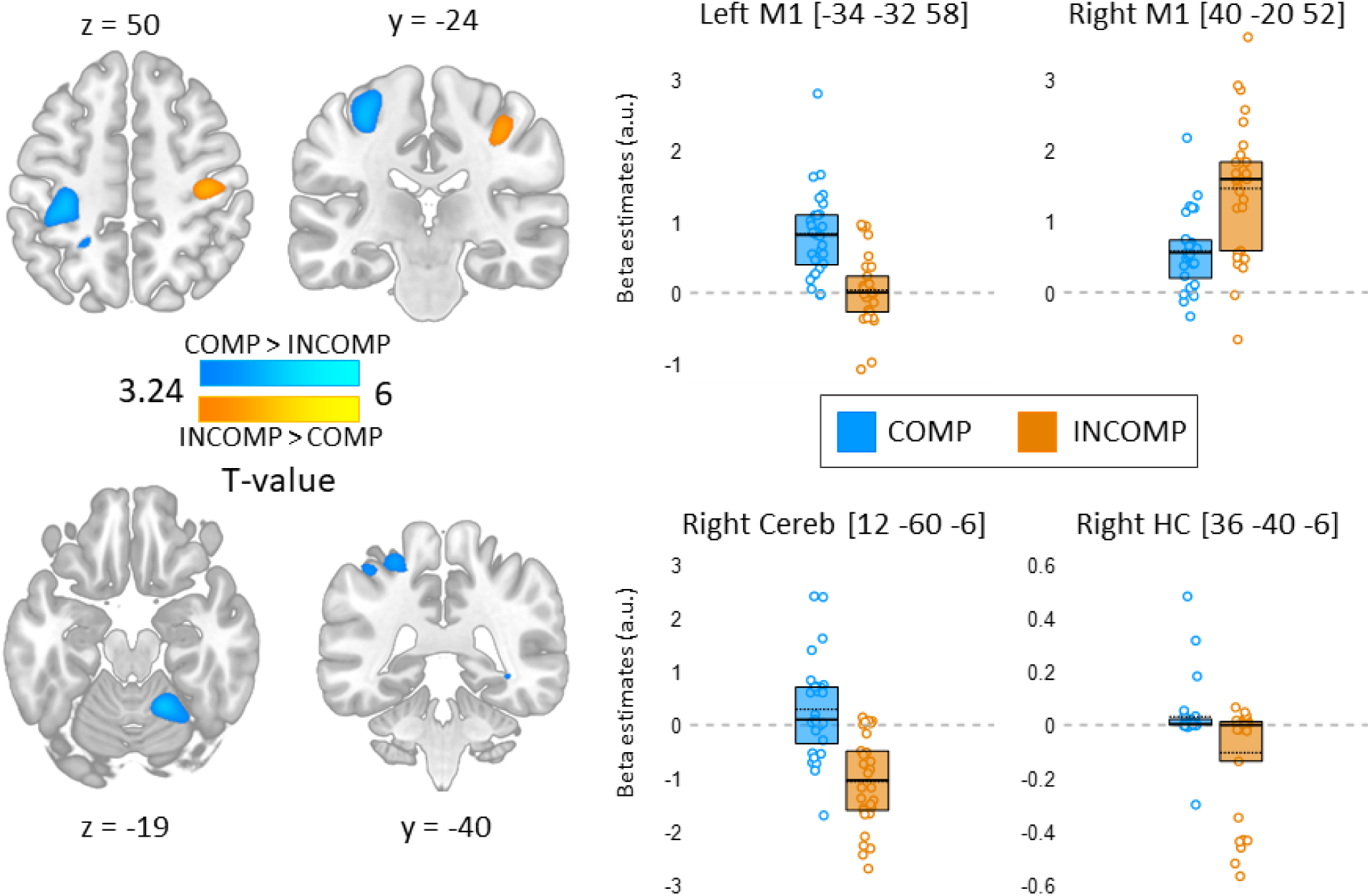
Left panel. Brain regions showing a significant difference in task-related brain activity between the COMP (N=29) and INCOMP (N=30) groups (blue: COMP > INCOMP and orange: INCOMP > COMP). Statistical images are thresholded at p<0.001 uncorrected. Results are presented in MNI152 space. **Right panel**. Beta values (arbitrary units, a.u.) for the peak coordinates are displayed for the COMP (blue) and INCOMP (orange) groups. Full horizontal bars indicate the median and dotted horizontal bars the mean. Colored circles represent individual data points. Boxes represent the interquartile range (IQR). Cereb=cerebellum; HC=hippocampus.

To further investigate the lateralization observed in M1 during performance of the schema-compatible versus incompatible sequence, we examined whether schema-compatibility affected the task-related functional *connectivity* of these brain regions. To do so, we performed psycho-physiological interaction analyses and tested whether the whole-brain connectivity of the M1 seeds described above (i.e., left and right M1) differed between COMP and INCOMP groups. This analysis revealed that the strength of the task-related functional connectivity between the left M1 seed and a set of regions including the left hippocampus, the right putamen, the left intra parietal sulcus and the left cerebellum was significantly lower in the COMP compared to the INCOMP group (Figure 4A and Table 3C). Reduced connectivity in the COMP versus INCOMP group was also observed between the right M1 seed and the right cerebellum during task practice as compared to rest (Figure 4B and Table 3D).

**Figure 4:**
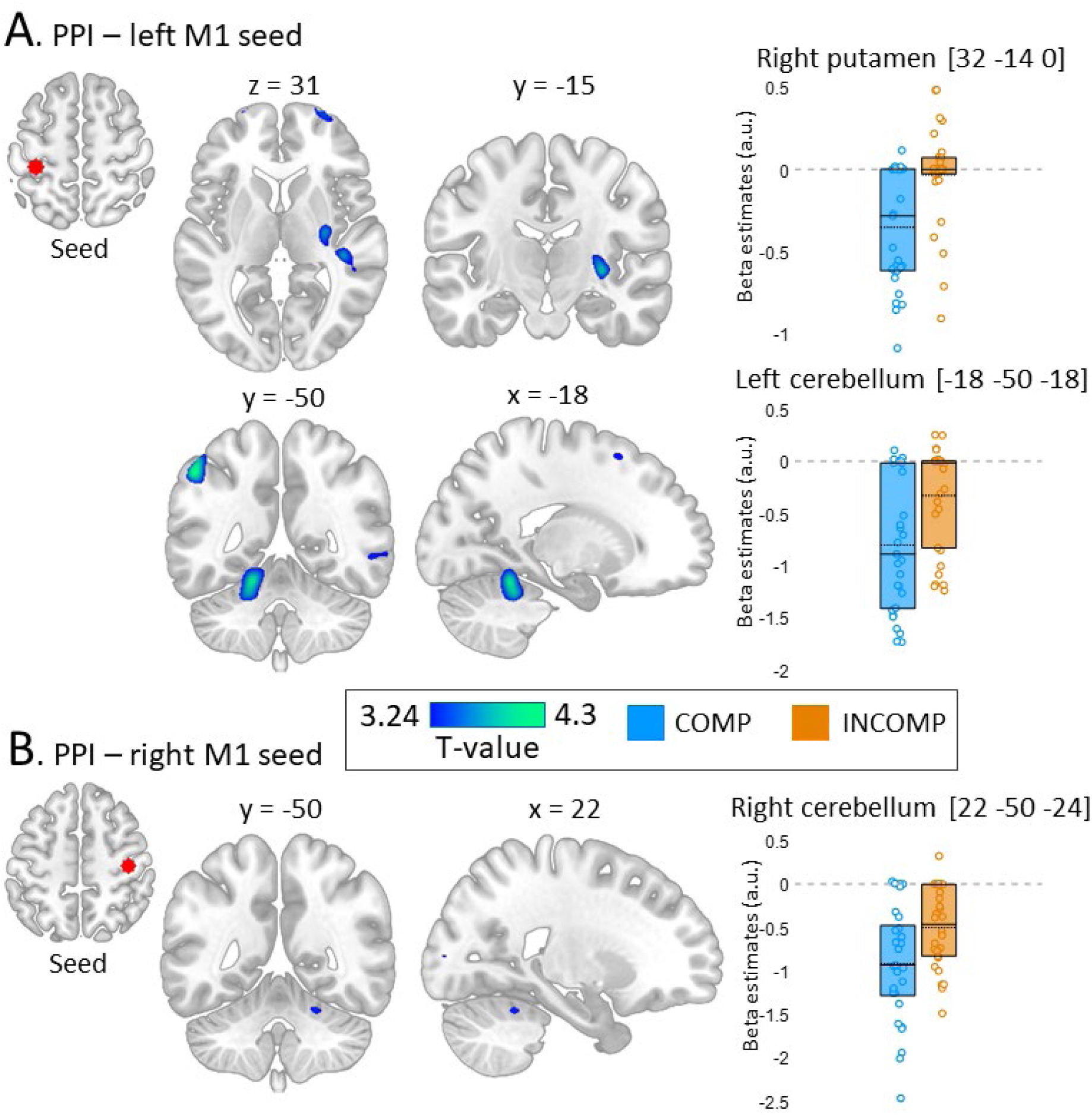
A. Left panel. Brain regions showing a significantly lower task-related functional connectivity with the left M1 (coordinate [-34 -32 58], seed region shown in red) in the COMP (N=29) versus INCOMP (N=30) group. **B. Left panel.** Brain regions showing a significantly lower task-related functional connectivity with the right M1 (coordinate [40 -20 52], seed region shown in red) in the COMP versus INCOMP group. In both panels, statistical images are thresholded at p<0.001 uncorrected. Results are presented in MNI152 space. **A-B. Right panels.** Beta values (arbitrary units, a.u.) for the peak coordinates are displayed for the COMP (blue) and INCOMP (orange) groups. Full horizontal bars indicate the median and dotted horizontal bars the mean. Colored circles represent individual data points. Boxes represent the interquartile range (IQR).

In sum, the results of the univariate MRI analyses suggest that the recruitment of the right M1 is greater during sequence practice in a schema-incompatible, compared to compatible, context, while a wider set of brain regions including the left M1, hippocampus and cerebellum is more activated in the schema-compatible, compared to incompatible, condition. Interestingly, both left and right M1 connectivity with striato-hippocampo-cerebellar regions was overall lower during practice in a compatible, compared to incompatible, context.

**Table 3:** Functional imaging results for Session 2 SRTT practice [between-group comparisons of task-related activity (a,b) and connectivity (c,d)].

| Area | x | y | z | k | T | $P_{corr}$ |
| --- | --- | --- | --- | --- | --- | --- |
| **A. Main effect of schema compatibility (COMP > INCOMP)** |  |  |  |  |  |  |
| R Lingual gyrus extending into R Cerebellar lobules IV-V | 12 | -60 | -6 | 455 | 6.08 | <0.003 |
| L Primary motor cortex | -34 | -32 | 58 | 406 | 5.23 | <0.003 |
| L Precuneus | -26 | -50 | 54 | 56 | 3.81 | 0.008 |
| L Supplementary motor area | -6 | -16 | 60 | 69 | 4.07 | 0.006 |
| R Hippocampus | 36 | -40 | -6 | 10 | 3.43 | 0.017 |
| R Cerebellar lobule VIII | 18 | -60 | -46 | 14 | 3.36 | 0.017 |
| **B. Main effect of schema incompatibility (INCOMP > COMP)** |  |  |  |  |  |  |
| R Primary motor cortex | 40 | -20 | 52 | 253 | 4.37 | 0.001 |
| **C. Psycho-physiological interaction, seed region left primary motor cortex (INCOMP > COMP)** |  |  |  |  |  |  |
| L Intra parietal sulcus | -50 | -54 | 52 | 62 | 4.61 | 0.005 |
| L Cerebellar lobules IV-V | -18 | -50 | -18 | 175 | 4.25 | 0.005 |
| L Cerebellar lobules IV-V | -20 | -48 | -24 | 175 | 4.12 | 0.005 |
| R Putamen | 32 | -14 | 0 | 34 | 4.04 | 0.006 |
| L Fusiform gyrus | -40 | -38 | -16 | 62 | 3.79 | 0.008 |
| L Hippocampus | -40 | -26 | -16 | 62 | 3.54 | 0.012 |
| R Primary sensory cortex | 30 | -30 | 58 | 87 | 3.75 | 0.008 |
| L Superior frontal cortex | -20 | 14 | 56 | 33 | 3.49 | 0.012 |
| **D. Psycho-physiological interaction, seed region right primary motor cortex (INCOMP > COMP)** |  |  |  |  |  |  |
| R Cerebellar lobule VI | 22 | -50 | -24 | 11 | 3.47 | 0.012 |

#### 3.2.2. Multivariate pattern analyses

The multivariate analyses of the MRI data examined group differences in brain patterns associated with each key/ordinal position pairing in the S2 sequences. Brain patterns were extracted from specific ROIs involved in motor learning, i.e., the hippocampus, putamen, left and right M1, premotor cortex and aSPL (note that all ROIs were bilateral except for M1, which was divided in left and right M1 based on the lateralization observed in the univariate analyses presented above). Exploratory analyses on two ROIs derived from the declarative memory schema literature (i.e., the medial prefrontal cortex and the angular gyrus) are presented in the Supplementary material (Table S7, Figure S1).

The results of the 2 *groups* (COMP/INCOMP) by 8 *ordinal positions* (1 through 8) ANOVAs - shown in Figure 5 and reported in Table 4 - reveal a significant main effect of ordinal position in all our examined ROIs. Data inspection suggests that this effect is driven by a pronounced edge effect whereby similarity values on the first and last sequence elements tended to be greater than on all other sequence elements across most of the ROIs (i.e., the average similarity of ordinal positions 1 and 8 was greater than that of ordinal positions 2-7 in all ROIs [paired-samples t-tests, all *p_s_*<0.01] except the right M1 [*p*=0.35]). These results are consistent with previous work demonstrating such effect across a large number of motor sequence learning-related ROIs (Dolfen et al., 2024). In contrast, no significant main effect of group was detected in any of the examined ROIs. Interestingly, significant interactions of the factors group and ordinal position were observed in both the left and right M1 as well as in the premotor cortex, suggesting that the representation of sequence elements in these ROIs – but not in the aSPL, hippocampus, and putamen – was influenced by schema-compatibility.

To better characterize the differences in neural representation of sequence elements between schema-compatible and -incompatible conditions, we performed follow-up pairwise analyses in the three regions showing a significant group x ordinal position interaction effect (results of follow-up analyses are presented in Table 5). In both the **left M1** and the **premotor cortex**, a significant group difference in similarity values was detected on the first sequence element such that pattern similarity was greater in the INCOMP, as compared to the COMP, group. These results suggest a greater representation of the first element of the sequence in these regions when practice takes place in a schema-incompatible, compared to a schema-compatible, context. Additionally, multivoxel activation patterns in **both left and right M1** showed increased similarity in the schema-compatible condition before, during and after the introduction of the 1^st^ *novel* key in the sequence (2^nd^, 3^rd^ and 4^th^ sequence elements), implicating M1 in the representation of novel sequence elements only when these are presented in a schema-compatible context. This effect was less pronounced in the **premotor cortex** as it was limited to the key following the novel element (4^th^ sequence element). No significant group effects were observed on the other sequence elements including around or on the 2^nd^ novel key.

Altogether, the multivoxel activation patterns in these motor cortical regions might reflect different processes related to (i) the salience of the ordinal mismatch on the first sequence element under incompatible condition, and (ii) the integration of (the first) novel element into a compatible schema.

**Figure 5:**
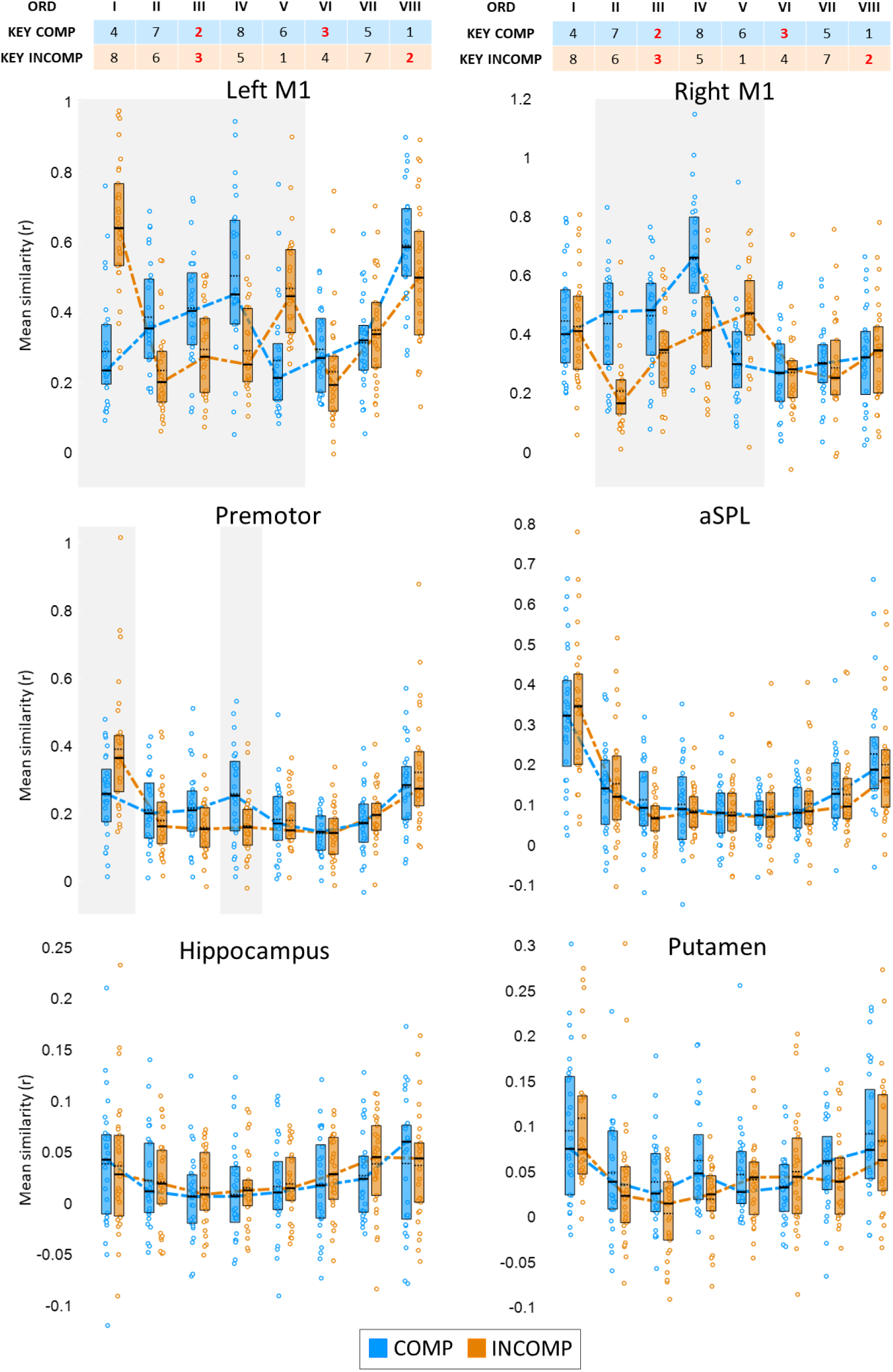
Mean pattern similarity per key/ordinal position pairing, for each experimental group and ROI. The top table represents the ordinal (ORD) position of each element in the sequence (position 1 to 8) sequence of movements (here, each number represents a finger with 1 and 8 corresponding to the left and right little fingers, respectively). Full horizontal bars indicate the median and dotted horizontal bars the mean. Coloured dashed lines connect the medians in each experimental group. Coloured circles represent individual data. Boxes represent the interquartile range (IQR). Shaded grey rectangles denote key/ordinal position pairings for which follow-up tests indicated a significant difference in similarity across groups (see Table 5). Premotor = premotor cortex; aSPL = anterior superior parietal lobule.

**Table 4:** Results of multivariate pattern analyses for the 2 groups (COMP/INCOMP) x 8 ordinal positions (Ord; 1 through 8) ANOVAs.

| ROI | Main effect Group |  |  |  | Main effect Ord |  |  |  | Group x Ord interaction |  |  |  |
| --- | --- | --- | --- | --- | --- | --- | --- | --- | --- | --- | --- | --- |
| | df | F | p<sub>corr</sub> | Part $\eta^2$ | df | F | p<sub>corr</sub> | Part $\eta^2$ | df | F | p<sub>corr</sub> | Part $\eta^2$ |
| L M1 | 1,57 | 0.04 | 0.93 | 0.001 | 4.21,239.86 | 32.19 | **<0.001\*** | 0.361 | 4.21,239.86 | 37.20 | **<0.002\*** | 0.395 |
| R M1 | 1,57 | 3.54 | 0.42 | 0.058 | 4.40,250.82 | 30.14 | **<0.001\*** | 0.346 | 4.40,250.82 | 17.14 | **<0.002\*** | 0.231 |
| PMC | 1,57 | 0.01 | 0.93 | 0.000 | 4.25,241.94 | 32.50 | **<0.001\*** | 0.363 | 4.25,241.94 | 10.00 | **<0.002\*** | 0.149 |
| aSPL | 1,57 | 0.17 | 0.93 | 0.003 | 3.90,222.00 | 18.32 | **<0.001\*** | 0.243 | 3.90,222.00 | 1.00 | 0.49 | 0.017 |
| HC | 1,57 | 0.30 | 0.93 | 0.005 | 3.98,226.86 | 3.99 | **0.004\*** | 0.065 | 3.98,226.86 | 0.28 | 0.89 | 0.005 |
| Put | 1,57 | 0.84 | 0.93 | 0.015 | 3.89,221.98 | 16.08 | **<0.001\*** | 0.220 | 3.89,221.98 | 2.35 | 0.09 | 0.040 |

L/R M1 = left/right primary motor cortex; PMC = premotor cortex; aSPL = anterior superior parietal lobule; HC = hippocampus; Put = putamen; df = degrees of freedom; part $\eta^2$ = partial eta squared. $P_{corr}$ indicates the p-value corrected for multiple comparisons (FDR correction for the number of ROIs). Significant p-values are marked with an asterisk and bold font.

**Table 5:** Results of independent samples t-tests (COMP vs INCOMP) for each of the 8 ordinal positions (Ord; 1 through 8).

| Ord | Left M1 |  |  | Right M1 |  |  | Premotor cortex |  |  |
| --- | --- | --- | --- | --- | --- | --- | --- | --- | --- |
|  | t | p<sub>corr</sub> | Cohen's d | t | p<sub>corr</sub> | Cohen's d | t | p<sub>corr</sub> | Cohen's d |
| I | -7.74 | **<0.002\*** | -2.02 | 0.39 | 0.80 | 0.10 | -3.22 | **0.01\*** | -0.84 |
| II | 4.08 | **<0.002\*** | 1.06 | 5.40 | **<0.004\*** | 1.41 | 1.16 | 0.42 | 0.30 |
| III | 3.11 | **0.005\*** | 0.81 | 2.97 | **0.01\*** | 0.77 | 2.24 | 0.08 | 0.58 |
| IV | 4.63 | **<0.002\*** | 1.22 | 4.93 | **<0.004\*** | 1.28 | 3.12 | **0.01\*** | 0.82 |
| V | -5.03 | **<0.002\*** | -1.31 | -3.07 | **0.01\*** | -0.80 | 0.12 | 0.94 | 0.03 |
| VI | 1.71 | 0.11 | 0.45 | 0.03 | 0.98 | 0.01 | -0.08 | 0.94 | -0.02 |
| VII | -0.96 | 0.34 | -0.25 | 0.48 | 0.80 | 0.12 | -0.97 | 0.45 | -0.25 |
| VIII | 1.93 | 0.08 | 0.50 | -0.50 | 0.80 | -0.13 | -1.13 | 0.42 | -0.29 |

Degrees of freedom = 57 for all tests with following exceptions: Left M1, ord IV: df=42.41; Right M1, ord II: df=52.20; Premotor cortex, ord IV: df=47.60. $P_{corr}$ indicates the p-value corrected for multiple comparisons (FDR correction for the number of sequence elements). Significant p-values are marked with an asterisk and bold font.

## 4. Discussion

In the present paper, we examined the neural processes underlying schema-mediated integration of novel information in motor memory. Compatibility with a previous cognitive-motor schema was examined with a protocol similar to our earlier research (King et al., 2019; Reverberi et al., 2023) in which the ordinal structure of the practiced sequence (i.e., the temporal position of movements in the motor sequence) was manipulated such that it was compatible or incompatible with that of the motor sequence learned on the previous day. Our behavioural results are in line with earlier observations (King et al., 2019) and suggest that performance on new sequential movements is enhanced in a context that is compatible with the previously acquired ordinal schema. Brain imaging results show that sequence practice in a schema-compatible (as compared to -incompatible) context resulted in an overall increase in left M1 activity and in a decrease in M1 connectivity with striato-hippocampo-cerebellar areas. Multivariate fMRI analyses indicated that multivoxel pattern similarity in both the left and right M1 as well as in the premotor cortex was increased around the first novel key in the sequence stream in a compatible compared to incompatible context. Altogether, these results suggest that motor cortical areas facilitate the integration of novel elements that are compatible with a pre-existing cognitive-motor schema.

Our behavioural results replicated those of our previous study (King et al., 2019), as they showed a performance advantage for the new sequence practiced in a context that was compatible, as compared to incompatible, with the ordinal framework learned the previous day. This group difference was observed during the test session prior to scanning but did not persist during the runs of task performed inside the MRI scanner. We speculate that the slower task pace in the MR scanner used to accommodate RSA might have attenuated these overall schema effect faster than in our previous research (King et al., 2019) in which both groups eventually reached the same level of performance after extensive practice. Altogether, the behavioral results confirm earlier observations of schema-mediated facilitation of the integration of new movements into motor memory (King et al., 2019).

Our activation-based univariate analyses revealed that sequence practice in a schema-compatible, versus incompatible, context resulted in greater task-related activity in a set of brain regions including the left M1, the left precuneus, left SMA, right hippocampus and right cerebellum. In contrast, practice in an incompatible context showed greater activation of the right M1. The schema-mediated lateralization effect in the primary motor cortex is intriguing as it cannot be attributed to differences in the nature of the movements performed during task practice, given that the two experimental groups performed the same bimanual sequence (only the starting points differed between groups). We suggest that these differences might instead reflect distinct cognitive processes. Previous literature has suggested a left hemispheric specialization in the learning of new motor skills and particularly motor sequences. Specifically, while right hemisphere recruitment has been shown to be limited to the execution of contralateral hand movements, the left hemisphere is engaged during the learning of new sequences performed with both the contralateral and the ipsilateral hands (Grafton et al., 1995, 1998, 2002). In line with these observations, left but not right hemisphere stroke has been associated with deficits in the encoding and generation of sequential movements (Harrington & Haaland, 1991; Kimura, 1977). Additionally, non-invasive brain stimulation studies showed that left M1 stimulation benefits motor learning more than sham or right M1 stimulation (Schambra et al., 2011). Thus, the heightened activity in the left M1 observed in the current study in a schema-compatible, compared to incompatible, context may reflect improved sequence learning when the ordinal framework is congruent with previous knowledge. Consistent with this interpretation, task practice in a schema-compatible group also recruited a set of brain regions known to play a critical role in motor learning processes, including the SMA and the parietal cortex (Albouy et al., 2008; Doyon et al., 2009; Fernández-Seara et al., 2009; Grafton et al., 2002; Hikosaka et al., 1996; Jacobacci et al., 2020; Penhune & Steele, 2012). In contrast, the recruitment of the right M1 in the schema-incompatible group might reflect increased use of control and attentional processes potentially induced by the ordinal incongruency. The right hemisphere in general is known to be critical for the control of visuospatial processing and spatial attention (Corballis, 2003; Coull & Nobre, 1998; Winstein & Pohl, 1995), and the primary motor cortex has also been shown to participate in attention to movement and motor imagery (Binkofski et al., 2002; Johansen-Berg & Matthews, 2002; Lotze et al., 1999; Porro et al., 1996; Tagaris et al., 1998; Tomasino et al., 2005, see Bhattacharjee et al., 2021 for a review). Together with the evidence reviewed above that the right M1 is less prone than the left M1 to support learning, greater right M1 activation during learning of a schema-incompatible sequence may reflect a need for increased attention to the visual stimuli when presented in an ordinal framework that is incompatible with previous experience.

Differences in activation between the compatible and incompatible groups were also observed in the cerebellum and in the hippocampus. Interestingly, these effects were driven by negative parameter estimates in the schema-incompatible group while estimates were around zero in the compatible group. These results suggest that the cerebellum and the hippocampus were more *deactivated* during task practice in the incompatible as compared to the compatible group. Our results therefore support the hypothesis that hippocampal responses are modulated by schema-compatibility in the motor memory domain but with a direction that was opposite to our expectations. Indeed, in the declarative memory domain, hippocampal involvement has been described to be *greater* during schema-incompatible as compared to compatible learning (van Kesteren et al., 2009, 2013). Additionally, recent neuroimaging work suggests that the hippocampus plays a critical role in the acquisition of new motor sequences (Albouy et al., 2013, 2015; Fernández-Seara et al., 2009; Jacobacci et al., 2020) and represents ordinal information about movements in a learned sequence (Dolfen et al., 2024). According to these previous studies, we hypothesized that the hippocampus would be involved in the learning of the schema-incompatible motor sequence but not in the learning of the schema-compatible sequence which was expected to be rapidly assimilated into cortical storage. We therefore predicted that the practice of the incompatible sequence would result in *increased* task-related hippocampal activity. In contrast to these expectations, the current results point to greater *deactivation* of the hippocampus in the schema-incompatible group. An alternative explanation for the observed results is that hippocampal activity was *increased* at *rest* as compared to *practice* in the schema-incompatible, compared to -compatible, group (see negative beta values in Figure 3). Recently, increased attention has been paid to periods of quiet rest interleaved with task practice during sequence learning as they are proposed to host fast offline motor memory consolidation (Bönstrup et al., 2019). Interestingly, these rest periods have been associated with hippocampal responses that are thought to reflect the reactivation of motor task-related patterns (Buch et al., 2021; Gann et al., 2023 Jacobacci et al., 2020). Based on this previous evidence, it is therefore tempting to speculate that the pattern of hippocampal results reflects increased hippocampal reactivations of the new – incompatible – motor sequence task during rest. This is however speculative and warrants further investigation into the reactivation of brain patterns elicited by schema-incompatible (vs. compatible) motor learning.

Our connectivity analyses consistently showed that both the left and right M1 were less connected to other task-relevant brain regions during sequence practice (as compared to rest) in a schema-compatible, as compared to incompatible, context. Specifically, the connectivity between the left M1 and a set of brain regions including motor cortical, striatal, hippocampal and cerebellar areas was decreased under schema-compatible conditions. An analogous pattern of reduced task-related connectivity was also observed in the compatible group between the right M1 and the right cerebellum. The connectivity results observed in the current study are generally in line with previous observations of decreased connectivity in schema-congruent conditions in the declarative memory domain. Specifically, previous research has reported decreased hippocampo-cortical (prefrontal) connectivity during the encoding of schema-compatible, compared to schema-incompatible, novel movie scenes (van Kesteren et al., 2010) and novel factual information (van Kesteren et al., 2014). These results are thought to reflect a decreased need for crosstalk between distant brain regions (in particular between the hippocampus and the prefrontal cortex) in schema-compatible conditions, whereby integration of novel information into memory occurs directly in the neocortex, effectively bypassing hippocampal involvement (van Kesteren et al., 2010). The current results extend these findings to the motor memory domain and suggest that schema-compatibility resulted in a decreased crosstalk between the motor cortex and distant task-relevant brain regions including the hippocampus, striatum, and cerebellum. In the motor domain, such connectivity decreases are usually observed as learning progresses (Tzvi et al., 2014) while initial motor learning is generally associated with increased inter- and intra-hemispheric functional coupling between the sensorimotor, premotor, and supplementary motor cortices (Sun et al., 2007) and between the hippocampus and the striatum (Albouy et al., 2013). These results suggest that schema-compatibility might have accelerated network segregation processes that are usually observed as a function of practice (Tzvi et al., 2014). As above, an alternative explanation for the observed results is that connectivity was *increased* at *rest* as compared to *practice* in the schema-compatible, compared to incompatible, group (see beta estimates in Figure 4). This alternative explanation is in line with previous studies reporting increased functional connectivity *at rest* in hippocampo-frontal networks after encoding of schema-compatible material (Liu et al., 2018; Schlichting & Preston, 2016; but see van Kesteren et al., 2010 for a different account). Such connectivity increase has been associated with greater coarseness of schema-compatible memories (Audrain & McAndrews, 2022), with the loss of detail suggesting increased consolidation and reliance on neocortical retrieval for schema-compatible information (Sekeres et al., 2018).

The results of our key-level multivariate pattern analyses revealed that, in the bilateral primary motor and premotor cortices, pattern similarity was significantly increased for those sequence elements neighboring the 1^st^ novel key in the sequence, but only when practiced in a schema-compatible, compared to incompatible context. The heightened representation of the 1^st^ novel key in the sequence (i.e., to the first element not belonging to the previously acquired schema), together with the increased left M1 recruitment observed during overall sequence practice in the compatible group, suggest a role for motor cortices in the rapid integration of novel schema-compatible information into motor memory. We speculate that M1 and the premotor cortex may play a role in the adaptation of the previously acquired sequence representation to accommodate new movements into memory if the new material to learn is compatible with previous experience. Previous studies using multivariate pattern analyses demonstrated that multivoxel patterns in M1 and the premotor cortex represent information about both individual finger movements (Berlot et al., 2020; Dolfen et al., 2024; Yokoi et al., 2018; Yokoi & Diedrichsen, 2019), as well as movement chunks and entire sequences (Kornysheva & Diedrichsen, 2014; Yokoi & Diedrichsen, 2019). Additionally, the premotor cortex is known to carry ordinal position information in streams of movements (Dolfen et al., 2024; Kornysheva & Diedrichsen, 2014) which may provide a temporal scaffold in which movements are integrated during task practice. We propose that in the current study, the specific responses of M1 and the premotor cortex around the 1^st^ novel key in the schema-compatible sequence may reflect these regions’ capacity to adapt the previously learned ordinal-based cognitive motor-schema (King et al., 2019) to assimilate the novel material.

The multivariate MRI analyses also show that when practice takes place in a schema-incompatible context, the left (but not the right) M1 and the premotor cortex displayed higher similarity for the 1^st^ element of the sequence. A first key effect was previously reported in M1, with M1 activation patterns largely depending on the sequence’s starting element (Yokoi et al., 2018). The current results extend these findings and show that this first key effect is (i) lateralized to the left M1, (ii) also observed in the premotor cortex, and (ii) modulated by previous experience as it is more pronounced when motor sequences are practiced in a context that is incompatible (as compared to compatible) with a previously acquired cognitive-motor schema. Consistent with the motor and premotor cortices’ sensitivity to novel keys in a compatible context discussed above, and considering that the schema-incompatible sequence, contrary to the compatible one, is different from the previously learned schema from the very 1^st^ keypress, the first key effect observed in our study may thus reflect an ordinal/key mapping mismatch. However, as no increase in similarity was observed in the schema-incompatible context after the first key, even though all other keys presented a similar ordinal mismatch, we speculate that this effect more likely reflects the salience of the ordinal mismatch at the start of the sequence. Indeed, in the incompatible group, no increases of pattern similarity were observed during or following performance of the novel keys. We argue that both the first key salience effect observed on the multivoxel patterns of the left M1 and the premotor cortex as well as the more general incompatibility effect observed on the amplitude of the BOLD signal in the right M1 may reflect increased attentional need during sequence performance in a schema-incompatible context.

In summary, our results indicate that schema-mediated motor learning is supported by increased activity in the left primary motor cortex and a decreased crosstalk between the motor cortex and distant task-relevant brain regions including the hippocampus, striatum, and cerebellum. Our findings also suggest that multivoxel activation patterns can be altered in the primary and the premotor cortices to assimilate new movements that are compatible with the previously acquired cognitive motor-schema. These results deepen our understanding of the neural processes underlying schema-mediated learning in the motor memory domain and suggest the involvement of motor cortical regions in schema-mediated integration of novel movements into memory.

## 5. Data and code availability

Raw data, analyzed data corresponding to the figures presented in the text, and the scripts used to produce them are publicly available on OSF (https://osf.io/xnrg8).

## 6. Author contributions

S. Reverberi: Methodology, Software, Formal analysis, Investigation, Data curation, Writing – original draft, Writing – review & editing, Visualization; N. Dolfen: Methodology, Software, Writing – review & editing; B.R. King: Conceptualization, Writing – review & editing, Funding acquisition; G. Albouy: Conceptualization, Writing – review & editing, Supervision, Project administration, Funding acquisition.

## References

Albouy, G., Fogel, S., King, B. R., Laventure, S., Benali, H., Karni, A., Carrier, J., Robertson, E. M., & Doyon, J. (2015). Maintaining vs. enhancing motor sequence memories: Respective roles of striatal and hippocampal systems. NeuroImage. 10.1016/j.neuroimage.2014.12.049

Albouy, G., King, B. R., Maquet, P., & Doyon, J. (2013). Hippocampus and striatum: Dynamics and interaction during acquisition and sleep-related motor sequence memory consolidation. In Hippocampus (Vol. 23, Issue 11, pp. 985–1004). 10.1002/hipo.22183

Albouy, G., Sterpenich, V., Balteau, E., Vandewalle, G., Desseilles, M., Dang-Vu, T., Darsaud, A., Ruby, P., Luppi, P. H., Degueldre, C., Peigneux, P., Luxen, A., & Maquet, P. (2008). Both the Hippocampus and Striatum Are Involved in Consolidation of Motor Sequence Memory. Neuron, 58(2), 261–272. 10.1016/J.NEURON.2008.02.008

Amunts, K., Mohlberg, H., Bludau, S., & Zilles, K. (2020). Julich-Brain: A 3D probabilistic atlas of the human brain’s cytoarchitecture. Science, 369(6506), 988–992. 10.1126/SCIENCE.ABB4588/SUPPL_FILE/ABB4588_AMUNTS_SM.PDF

Atienza, M., Crespo-Garcia, M., & Cantero, J. L. (2011). Semantic Congruence Enhances Memory of Episodic Associations: Role of Theta Oscillations. Journal of Cognitive Neuroscience, 23(1), 75–90. 10.1162/JOCN.2009.21358

Audrain, S., & McAndrews, M. P. (2022). Schemas provide a scaffold for neocortical integration of new memories over time. Nature Communications 2022 13: 1, *13*(1), 1–16. 10.1038/s41467-022-33517-0

Beck, A. T., Epstein, N., Brown, G., & Steer, R. A. (1988). An Inventory for Measuring Clinical Anxiety: Psychometric Properties. Journal of Consulting and Clinical Psychology, 56(6), 893–897. 10.1037/0022-006X.56.6.893

Beck, A. T., Steer, R. A., Ball, R., & Ranieri, W. F. (1996). Comparison of Beck Depression Inventories-IA and-II in Psychiatric Outpatients. Journal of Personality Assessment, 67(3), 588–597. 10.1207/S15327752JPA6703_13

Bein, O., Livneh, N., Reggev, N., Gilead, M., Goshen-Gottstein, Y., & Maril, A. (2015). Delineating the Effect of Semantic Congruency on Episodic Memory: The Role of Integration and Relatedness. PLOS ONE, 10(2), e0115624. 10.1371/JOURNAL.PONE.0115624

Benjamini, Y., & Hochberg, Y. (1995). Controlling the False Discovery Rate: A Practical and Powerful Approach to Multiple Testing. Journal of the Royal Statistical Society: Series B (Methodological), 57(1), 289–300. 10.1111/J.2517-6161.1995.TB02031.X

Berlot, E., Popp, N. J., & Diedrichsen, J. (2020). A critical re-evaluation of fmri signatures of motor sequence learning. ELife, 9, 1–24. 10.7554/ELIFE.55241

Bhattacharjee, S., Kashyap, R., Abualait, T., Annabel Chen, S. H., Yoo, W. K., & Bashir, S. (2021). The Role of Primary Motor Cortex: More Than Movement Execution. Journal of Motor Behaviour, 53(2), 258–274. 10.1080/00222895.2020.1738992

Binkofski, F., Fink, G. R., Geyer, S., Buccino, G., Gruber, O., Shah, N. J., Taylor, J. G., Seitz, R. J., Zilles, K., & Freund, H. J. (2002). Neural activity in human primary motor cortex areas 4a and 4p is modulated differentially by attention to action. Journal of Neurophysiology, 88(1), 514–519. 10.1152/JN.2002.88.1.514/ASSET/IMAGES/LARGE/9K0722431102.JPEG

Bonasia, K., Sekeres, M. J., Gilboa, A., Grady, C. L., Winocur, G., & Moscovitch, M. (2018). Prior knowledge modulates the neural substrates of encoding and retrieving naturalistic events at short and long delays. Neurobiology of Learning and Memory, 153, 26–39. 10.1016/J.NLM.2018.02.017

Bönstrup, M., Iturrate, I., Thompson, R., Cruciani, G., Censor, N., & Cohen, L. G. (2019). A Rapid Form of Offline Consolidation in Skill Learning. Current Biology : CB, 29(8), 1346–1351.e4. 10.1016/J.CUB.2019.02.049

Buch, E. R., Claudino, L., Quentin, R., Bönstrup, M., & Cohen, L. G. (2021). Consolidation of human skill linked to waking hippocampo-neocortical replay. Cell Reports, 35(10), 109193. 10.1016/J.CELREP.2021.109193

Buysse, D. J., Reynolds, C. F., Monk, T. H., Berman, S. R., & Kupfer, D. J. (1989). The Pittsburgh sleep quality index: A new instrument for psychiatric practice and research. Psychiatry Research, 28(2), 193–213. 10.1016/0165-1781(89)90047-4

Corballis, P. M. (2003). Visuospatial processing and the right-hemisphere interpreter. Brain and Cognition, 53(2), 171–176. 10.1016/S0278-2626(03)00103-9

Coull, J. T., & Nobre, A. C. (1998). Where and When to Pay Attention: The Neural Systems for Directing Attention to Spatial Locations and to Time Intervals as Revealed by Both PET and fMRI. Journal of Neuroscience, 18(18), 7426–7435. 10.1523/JNEUROSCI.18-18-07426.1998

Cycowicz, Y. M., Nessler, D., Horton, C., & Friedman, D. (2008). Retrieving object color: The influence of color congruity and test format. NeuroReport, 19(14), 1387–1390. 10.1097/WNR.0B013E32830C8DF1

Dayan, E., & Cohen, L. G. (2011). Neuroplasticity Subserving Motor Skill Learning. Neuron, 72(3), 443–454. 10.1016/J.NEURON.2011.10.008

Dinges, D. F., & Powell, J. W. (1985). Microcomputer analyses of performance on a portable, simple visual RT task during sustained operations. In Behaviour Research Methods, Instruments, & Computers (Vol. 17, Issue 6).

Dolfen, N., Reverberi, S., beeck, H. Op de, King, B. R., & Albouy, G. (2024). The hippocampus represents information about movements in their temporal position in a learned motor sequence. *Journal of Neuroscience*, e0584242024. 10.1523/JNEUROSCI.0584-24.2024

Doyon, J., Bellec, P., Amsel, R., Penhune, V., Monchi, O., Carrier, J., Lehéricy, S., & Benali, H. (2009). Contributions of the basal ganglia and functionally related brain structures to motor learning. Behavioural Brain Research, 199(1), 61–75. 10.1016/J.BBR.2008.11.012

Doyon, J., Penhune, V., & Ungerleider, L. G. (2003). Distinct contribution of the cortico-striatal and cortico-cerebellar systems to motor skill learning. Neuropsychologia, 41(3), 252–262. 10.1016/S0028-3932(02)00158-6

Durrant, S. J., Cairney, S. A., McDermott, C., & Lewis, P. A. (2015). Schema-conformant memories are preferentially consolidated during REM sleep. Neurobiology of Learning and Memory, 122, 41–50. 10.1016/j.nlm.2015.02.011

Ellis, B. W., Johns, M. W., Lancaster, R., Raptopoulos, P., Angelopoulos, N., & Priest, R. G. (1981). The St. Mary’s Hospital sleep questionnaire: A study of reliability. Sleep, 4(1), 93–97. 10.1093/SLEEP/4.1.93

Fan, L., Li, H., Zhuo, J., Zhang, Y., Wang, J., Chen, L., Yang, Z., Chu, C., Xie, S., Laird, A. R., Fox, P. T., Eickhoff, S. B., Yu, C., & Jiang, T. (2016). The Human Brainnetome Atlas: A New Brain Atlas Based on Connectional Architecture. *Cerebral Cortex (New York*, NY), 26(8), 3508. 10.1093/CERCOR/BHW157

Faul, F., Erdfelder, E., Lang, A. G., & Buchner, A. (2007). G*Power 3: a flexible statistical power analysis program for the social, behavioural, and biomedical sciences. Behaviour Research Methods, 39(2), 175–191. 10.3758/BF03193146

Fernández-Seara, M. A., Aznárez-Sanado, M., Mengual, E., Loayza, F. R., & Pastor, M. A. (2009). Continuous performance of a novel motor sequence leads to highly correlated striatal and hippocampal perfusion increases. NeuroImage, 47(4), 1797–1808. 10.1016/J.NEUROIMAGE.2009.05.061

Gann, M. A., Dolfen, N., King, B. R., Robertson, E. M., & Albouy, G. (2023). Prefrontal stimulation as a tool to disrupt hippocampal and striatal reactivations underlying fast motor memory consolidation. Brain Stimulation, 16(5), 1336–1345.

Gitelman, D. R., Penny, W. D., Ashburner, J., & Friston, K. J. (2003). Modeling regional and psychophysiologic interactions in fMRI: the importance of hemodynamic deconvolution. NeuroImage, 19(1), 200–207. 10.1016/S1053-8119(03)00058-2

Grafton, S. T., Hazeltine, E., & Ivry, R. (1995). Functional mapping of sequence learning in normal humans. Journal of Cognitive Neuroscience, 7(4), 497–510. 10.1162/JOCN.1995.7.4.497

Grafton, S. T., Hazeltine, E., & Ivry, R. B. (1998). Abstract and Effector-Specific Representations of Motor Sequences Identified with PET. Journal of Neuroscience, 18(22), 9420–9428. 10.1523/JNEUROSCI.18-22-09420.1998

Grafton, S. T., Hazeltine, E., & Ivry, R. B. (2002). Motor sequence learning with the nondominant left hand: A PET functional imaging study. Experimental Brain Research, 146(3), 369–378. 10.1007/S00221-002-1181-Y/METRICS

Harrington, D. L., & Haaland, K. Y. (1991). Hemispheric specialization for motor sequencing: Abnormalities in levels of programming. Neuropsychologia, 29(2), 147–163. 10.1016/0028-3932(91)90017-3

Haxby, J. V., Gobbini, M. I., Furey, M. L., Ishai, A., Schouten, J. L., & Pietrini, P. (2001). Distributed and overlapping representations of faces and objects in ventral temporal cortex. *Science (New York*, N.Y.), 293(5539), 2425– 2430. 10.1126/SCIENCE.1063736

Heikkilä, J., Alho, K., Hyvönen, H., & Tiippana, K. (2015). Audiovisual semantic congruency during encoding enhances memory performance. Experimental Psychology, 62(2), 123–130. 10.1027/1618-369/A000279

Hikosaka, O., Sakai, K., Miyauchi, S., Takino, R., Sasaki, Y., & Pütz, B. (1996). Activation of human presupplementary motor area in learning of sequential procedures: a functional MRI study. 10.1152/Jn.1996.76.1.617, 76(1), 617–621. 10.1152/JN.1996.76.1.617

Horne, J. A., & Ostberg, O. (1976). A self-assessment questionnaire to determine morningness-eveningness in human circadian rhythms. International Journal of Chronobiology, 4(2), 97–110. https://europepmc.org/article/med/1027738

Jacobacci, F., Armony, J. L., Yeffal, A., Lerner, G., Amaro, E., Jovicich, J., Doyone, J., & Della-Maggiore, V. (2020). Rapid hippocampal plasticity supports motor sequence learning. Proceedings of the National Academy of Sciences of the United States of America, 117(38), 23898–23903. 10.1073/PNAS.2009576117/SUPPL_FILE/PNAS.2009576117.SAPP.PDF

Johansen-Berg, H., & Matthews, P. M. (2002). Attention to movement modulates activity in sensori-motor areas, including primary motor cortex. Experimental Brain Research, 142(1), 13–24. 10.1007/S00221-001-0905-8/METRICS

J. P. Verma. (2015). Repeated measures design for empirical researchers (Wiley, Ed.).

Keidel, J. L., Oedekoven, C. S. H., Tut, A. C., & Bird, C. M. (2018). Multiscale Integration of Contextual Information During a Naturalistic Task. Cerebral Cortex, 28(10), 3531–3539. 10.1093/CERCOR/BHX218

Kimura, D. (1977). Acquisition of a motor skill after left-hemisphere damage. Brain : A Journal of Neurology, 100(3), 527–542. 10.1093/BRAIN/100.3.527

King, B. R., Dolfen, N., Gann, M. A., Renard, Z., Swinnen, S. P., & Albouy, G. (2019). Schema and Motor-Memory Consolidation. Psychological Science, 30(7), 963–978. 10.1177/0956797619847164

Kleiner, M., Brainard, D., Pelli, D., Ingling, A., Murray, R., & Broussard, C. (2007). What’s new in psychtoolbox-3. Perception, 36(14), 1–16. https://nyuscholars.nyu.edu/en/publications/whats-new-in-psychtoolbox-3

Kornysheva, K., & Diedrichsen, J. (2014). Human premotor areas parse sequences into their spatial and temporal features. ELife, 3, e03043. 10.7554/ELIFE.03043

Lewis, P. A., & Durrant, S. J. (2011). Overlapping memory replay during sleep builds cognitive schemata. In Trends in Cognitive Sciences (Vol. 15, Issue 8, pp. 343–351). 10.1016/j.tics.2011.06.004

Liu, Z. X., Grady, C., & Moscovitch, M. (2018). The effect of prior knowledge on post-encoding brain connectivity and its relation to subsequent memory. NeuroImage, 167, 211–223. 10.1016/J.NEUROIMAGE.2017.11.032

Lotze, M., Montoya, P., Erb, M., Hülsmann, E., Flor, H., Klose, U., Birbaumer, N., & Grodd, W. (1999). Activation of cortical and cerebellar motor areas during executed and imagined hand movements: An fMRI study. Journal of Cognitive Neuroscience, 11(5), 491–501. 10.1162/089892999563553

Mur, M., Bandettini, P. A., & Kriegeskorte, N. (2009). Revealing representational content with pattern-information fMRI—an introductory guide. Social cognitive and affective neuroscience, 4(1), 101–109.

Nissen, M. J., & Bullemer, P. (1987). Attentional requirements of learning: Evidence from performance measures. Cognitive Psychology, 19(1), 1–32. 10.1016/0010-0285(87)90002-8

Oldfield, R. C. (1971). The assessment and analysis of handedness: the Edinburgh inventory. Neuropsychologia, 9(1), 97–113.

Oosterhof, N. N., Connolly, A. C., & Haxby, J. V. (2016). CoSMoMVPA: multi-modal multivariate pattern analysis of neuroimaging data in Matlab/GNU Octave. Frontiers in neuroinformatics, 10, 27.

Pan, S. C., & Rickard, T. C. (2015). Sleep and motor learning: Is there room for consolidation? Psychological Bulletin, 141(4), 812–834. 10.1037/BUL0000009

Patenaude, B., Smith, S. M., Kennedy, D. N., & Jenkinson, M. (2011). A Bayesian model of shape and appearance for subcortical brain segmentation. NeuroImage, 56(3), 907–922. 10.1016/J.NEUROIMAGE.2011.02.046

Penhune, V. B., & Steele, C. J. (2012). Parallel contributions of cerebellar, striatal and M1 mechanisms to motor sequence learning. Behavioural Brain Research, 226(2), 579–591. 10.1016/J.BBR.2011.09.044

Poldrack, R. A. (2007). Region of interest analysis for fMRI. Social Cognitive and Affective Neuroscience, 2(1), 67–70. 10.1093/SCAN/NSM006

Porro, C. A., Francescato, M. P., Cettolo, V., Diamond, M. E., Baraldi, P., Zuiani, C., Bazzocchi, M., & Di Prampero, P. E. (1996). Primary Motor and Sensory Cortex Activation during Motor Performance and Motor Imagery: A Functional Magnetic Resonance Imaging Study. Journal of Neuroscience, 16(23), 7688–7698. 10.1523/JNEUROSCI.16-23-07688.1996

Povilas Karvelis, & oyvindlr. (2024). povilaskarvelis/DataViz: v3.2.4 (v3.2.4). Zenodo. 10.5281/zenodo.12749045

Reverberi, S., Dolfen, N., Van Roy, A., Albouy, G., & King, B. R. (2023). Sleep does not influence schema-facilitated motor memory consolidation. PLoS One, 18(1), e0280591.

Schambra, H. M., Abe, M., Luckenbaugh, D. A., Reis, J., Krakauer, J. W., & Cohen, L. G. (2011). Probing for hemispheric specialization for motor skill learning: A transcranial direct current stimulation study. Journal of Neurophysiology, 106(2), 652–661. 10.1152/JN.00210.2011/ASSET/IMAGES/LARGE/Z9K0081108790005.JPEG

Schlichting, M. L., & Preston, A. R. (2016). Hippocampal–medial prefrontal circuit supports memory updating during learning and post-encoding rest. Neurobiology of Learning and Memory, 134(Part A), 91–106. 10.1016/J.NLM.2015.11.005

Sekeres, M. J., Schomaker, J., Nadel, L., & Tse, D. (2024). To update or to create? The influence of novelty and prior knowledge on memory networks. Philosophical Transactions B, 379(1906). 10.1098/RSTB.2023.0238

Sekeres, M. J., Winocur, G., & Moscovitch, M. (2018). The hippocampus and related neocortical structures in memory transformation. Neuroscience Letters, 680, 39–53. 10.1016/J.NEULET.2018.05.006

Squire, L. R., & Alvarez, P. (1995). Retrograde amnesia and memory consolidation: a neurobiological perspective. Current opinion in neurobiology, 5(2), 169–177.

Sun, F. T., Miller, L. M., Rao, A. A., & D’Esposito, M. (2007). Functional Connectivity of Cortical Networks Involved in Bimanual Motor Sequence Learning. Cerebral Cortex, 17(5), 1227–1234. 10.1093/CERCOR/BHL033

Tagaris, G. A., Richter, W., Kim, S. G., Pellizzer, G., Andersen, P., Uǧurbil, K., & Georgopoulos, A. P. (1998). Functional magnetic resonance imaging of mental rotation and memory scanning: a multidimensional scaling analysis of brain activation patterns. Brain Research Reviews, 26(2–3), 106–112. 10.1016/S0165-0173(97)00060-X

Tomasino, B., Borroni, P., Isaja, A., & Rumiati, R. I. (2005). The role of the primary motor cortex in mental rotation: a TMS study. Cognitive Neuropsychology, 22(3–4), 348–363. 10.1080/02643290442000185

Tse, D., Langston, R. F., Kakeyama, M., Bethus, I., Spooner, P. A., Wood, E. R., Witter, M. P., & Morris, R. G. M. (2007). Schemas and memory consolidation. Science, 316(5821), 76–82. 10.1126/science.1135935

Tse, D., Takeuchi, T., Kakeyama, M., Kajii, Y., Okuno, H., Tohyama, C., Bito, H., & Morris, R. G. M. (2011). Schema-dependent gene activation and memory encoding in neocortex. Science, 333(6044), 891–895. 10.1126/SCIENCE.1205274/SUPPL_FILE/TSE.SOM.PDF

Tzvi, E., Münte, T. F., & Krämer, U. M. (2014). Delineating the cortico-striatal-cerebellar network in implicit motor sequence learning. NeuroImage, 94, 222–230. 10.1016/J.NEUROIMAGE.2014.03.004

van Kesteren, M. T. R., Beul, S. F., Takashima, A., Henson, R. N., Ruiter, D. J., & Fernández, G. (2013). Differential roles for medial prefrontal and medial temporal cortices in schema-dependent encoding: From congruent to incongruent. Neuropsychologia, 51(12), 2352–2359. 10.1016/J.NEUROPSYCHOLOGIA.2013.05.027

van Kesteren, M. T. R., Fernández, G., Norris, D. G., & Hermans, E. J. (2009). Schema-dependent hippocampo-prefrontal connectivity during memory encoding and post-encoding rest. NeuroImage, 47, S54. 10.1016/S1053-8119(09)70170-3

van Kesteren, M. T. R., Fernández, G., Norris, D. G., & Hermans, E. J. (2010). Persistent schema-dependent hippocampal-neocortical connectivity during memory encoding and postencoding rest in humans. Proceedings of the National Academy of Sciences of the United States of America, 107(16), 7550–7555. 10.1073/PNAS.0914892107/SUPPL_FILE/PNAS.200914892SI.PDF

van Kesteren, M. T. R., Rijpkema, M., Ruiter, D. J., Morris, R. G. M., & Fernández, G. (2014). Building on Prior Knowledge: Schema-dependent Encoding Processes Relate to Academic Performance. Journal of Cognitive Neuroscience, 26(10), 2250–2261. 10.1162/JOCN_A_00630

van Kesteren, M. T. R., Ruiter, D. J., Fernández, G., & Henson, R. N. (2012). How schema and novelty augment memory formation. Trends in Neurosciences, 35(4), 211–219. 10.1016/j.tins.2012.02.001

Wagner, I. C., van Buuren, M., Kroes, M. C. W., Gutteling, T. P., van der Linden, M., Morris, R. G., & Fernández, G. (2015). Schematic memory components converge within angular gyrus during retrieval. ELife, 4(NOVEMBER2015). 10.7554/ELIFE.09668

Winstein, C. J., & Pohl, P. S. (1995). Effects of unilateral brain damage on the control of goal-directed hand movements. Experimental Brain Research, 105(1), 163–174. 10.1007/BF00242191/METRICS

Yokoi, A., Arbuckle, S. A., & Diedrichsen, J. (2018). The role of human primary motor cortex in the production of skilled finger sequences. Journal of Neuroscience, 38(6), 1430–1442. 10.1523/JNEUROSCI.2798-17.2017

Yokoi, A., & Diedrichsen, J. (2019). Neural Organization of Hierarchical Motor Sequence Representations in the Human Neocortex. Neuron, 103(6), 1178–1190.e7. 10.1016/J.NEURON.2019.06.017

