## Supplementary material for "Motor cortical areas facilitate schema-mediated integration of new motor information into memory"

#### Participant characteristics

Participant characteristics, vigilance, sleep duration and quality for the nights preceding the experimental sessions are reported in Table S1 and did not differ between experimental groups.

*Table S1: Participant demographics, sleep and vigilance data for each experimental group.*

| | **Group mean (SD)** | | | | |
|---|---|---|---|---|---|
| **Variable** | **COMP** | **INCOMP** | **t** | **p** | **Cohen's d** |
| N | 30 | 30 | | | |
| Female (n) | 20 | 20 | | | |
| Age (years) | 23.1 (2.8) | 23.5 (2.7) | -0.52 | 0.61 | -0.134 |
| BAI score | 3.0 (3.1) | 3.2 (3.3) | -0.24 | 0.81 | -0.063 |
| BDI score | 4.3 (3.8) | 3.6 (3.9) | 0.70 | 0.49 | 0.181 |
| Handedness score | 80.8 (14.6) | 84.0 (16.0) | -0.80 | 0.43 | -0.207 |
| PSQI score | 3.3 (1.8) | 2.9 (1.3) | 1.16 | 0.25 | 0.298 |
| Chronotype score | 50.6 (7.0) | 50.9 (7.0) | -0.20 | 0.84 | -0.052 |
| Average sleep duration, 3 nights prior S1 (hours) | 8:23 (0:44) | 8:22 (0:46) | 0.10 | 0.92 | 0.025 |
| Sleep duration, night prior S2 (hours) | 8:38 (0:58) | 8:38 (0:52) | 0.00 | >0.99 | 0.000 |
| Session 1 SMS duration (hours) | 8:09 (1:04) | 8:13 (0:41) | -0.27 | 0.79 | -0.069 |
| Session 2 SMS duration (hours) | 8:22 (0:56) | 8:11 (0:48) | 0.82 | 0.42 | 0.212 |
| Session 1 SMS quality | 4.0 (0.6) | 4.0 (0.7) | 0.10 | 0.92 | 0.025 |
| Session 2 SMS quality | 4.0 (0.5) | 4.1 (0.6) | -0.68 | 0.50 | -0.175 |
| Session 1 PVT (ms) | 298 (21) | 287 (30) | 1.73 | 0.09 | 0.446 |
| Session 2 PVT (ms) | 302 (25) | 290 (30) | 1.81 | 0.08 | 0.467 |

*Group means are presented, with standard deviation in parentheses. Average sleep duration for the 3 nights prior to S1 was assessed via self-reported sleep diaries. Sleep duration for the night prior to S2 was assessed via visual inspection of actigraphy data extracted from wrist-mounted device. Statistical differences between groups were assessed with independent samples t-tests. BAI = Beck Anxiety Inventory (Beck et al., 1988); BDI = Beck Depression Inventory (Beck et al., 1996); PSQI = Pittsburgh Sleep Quality Index (Buysse et al., 1989); SMS = St. Mary's Hospital Sleep Questionnaire (Ellis et al., 1981). Degrees of freedom=56 for Average sleep duration, 3 nights prior to S1, and for Sleep duration, night prior to S2; degrees of freedom=58 for all other variables.*

#### Random SRTT performance

*Table S2: Output of the statistical analyses assessing group differences in performance on the pseudo-random SRT task administered prior to the sequential SRT task in Session 1.*

| Effect | df | F | p | Partial $\eta^2$ |
|---|---|---|---|---|
| **A. Response Time** | | | | |
| Block | 2.83,163.99 | 10.08 | **<0.001*** | 0.148 |
| Block x Group | 2.83,163.99 | 0.48 | 0.69 | 0.008 |
| Group | 1,58 | 0.27 | 0.61 | 0.005 |
| **B. Accuracy** | | | | |
| Block | 3,174 | 1.11 | 0.35 | 0.019 |
| Block x Group | 3,174 | 0.18 | 0.91 | 0.003 |
| Group | 1,58 | 0.06 | 0.82 | 0.001 |

*Separate 4 (Block) by 2 (Group) ANOVAs were run per each variable (A: Response Time; B: Accuracy). The significant effect of block on RT reflects increased task familiarization with practice. The lack of group effect or block x group interaction indicate that general motor execution did not differ between experimental groups at baseline. Df=degrees of freedom. Significant p-values are marked with an asterisk and bold font.*

#### Small volume correction for univariate MRI analyses

*Table S3: Coordinates used for small volume correction for the data presented in Table 3 of the main text.*

| Area | x | y | z | Reference coordinate |
|---|---|---|---|---|
| R Lingual gyrus extending into R Cerebellar lobules IV-V | ±12 | -58 | -10 | (Destrebecqz et al., 2005) |
| L Primary motor cortex | ±36.2±3.0 | -22.3 ±4.6 | 57.0±6.1 | (Lehéricy et al., 2006) |
| L Precuneus | ±20 | -52 | 50 | (Fischer et al., 2005) |
| L Supplementary motor area | ±10 | -22 | 58 | (Penhune & Doyon, 2005) |
| R Hippocampus | ±32 | -42 | -4 | (Albouy et al., 2008) |
| R Cerebellar lobule VIII | ±22 | -62 | -48 | (Fischer et al., 2005) |
| R Primary motor cortex | ±36.2±3.0 | -22.3 ±4.6 | 57.0±6.1 | (Lehéricy et al., 2006) |
| L Intra parietal sulcus | ±48 | -50 | 60 | (Albouy et al., 2008) |
| L Cerebellar lobules IV-V | -18 | -44 | -18 | (Fischer et al., 2005) |
| L Cerebellar lobules IV-V | -18 | -44 | -18 | (Fischer et al., 2005) |
| R Putamen | 28 | -14 | -8 | (Albouy et al., 2008) |
| L Fusiform gyrus | ±42 | -34 | -12 | (Albouy et al., 2008) |
| L Hippocampus | ±42 | -34 | -12 | (Albouy et al., 2008) |
| R Primary sensory cortex | ±36.2±3.0 | -22.3 ±4.6 | 57.0±6.1 | (Lehéricy et al., 2006) |
| L Superior frontal cortex | ±22 | 12 | 54 | (Penhune & Doyon, 2002) |
| R Cerebellar lobule VI | 28 | -54 | -28 | (Dolfen et al., 2021) |
| L Anterior hippocampus | -18 | -16 | -14 | (Strange et al., 1999) |

#### ROI size for multivariate MRI analyses

*Table S4: Average number of voxels per each ROI used in representational similarity analyses per experimental group.*

| | Group mean (SD) | | | | |
|---|---|---|---|---|---|
| **ROI** | **COMP** | **INCOMP** | **t** | **p** | **Cohen's d** |
| Hippocampus | 877 (103) | 874 (85) | 0.12 | 0.91 | 0.031 |
| Putamen | 999 (135) | 1007 (112) | -0.27 | 0.79 | -0.070 |
| Left M1 | 782 (89) | 792 (91) | -0.42 | 0.68 | -0.110 |
| Right M1 | 671 (80) | 689 (74) | -0.89 | 0.38 | -0.232 |
| Premotor cortex | 1278 (155) | 1285 (154) | -0.18 | 0.86 | -0.046 |
| aSPL | 710 (105) | 727 (86) | -0.68 | 0.50 | -0.177 |
| Angular gyrus | 3842 (502) | 3822 (361) | 0.18 | 0.86 | 0.047 |
| mPFC | 1585 (197) | 1563 (183) | 0.46 | 0.65 | 0.119 |

*Group means are presented, with standard deviation in parentheses. Statistical differences between groups were assessed with independent samples t-tests. Degrees of freedom=57 for all tests. The angular gyrus and mPFC ROIs were considered exploratory analyses.*

#### Neuroimaging results: univariate analyses

##### Task practice > Rest contrast across both experimental groups

We examined which brain regions were recruited during task practice, compared to rest, across both experimental groups. As expected, task practice recruited a wide bilateral network of subcortical (putamen), cortical (M1, supplementary motor area (SMA), superior parietal lobule) and cerebellar regions across the two experimental groups which is in line with previous literature (Dayan & Cohen, 2011; Doyon et al., 2003). The corresponding results are reported in Table S5.

*Table S5: Functional imaging results for Session 2 SRTT practice versus rest, across both experimental groups.*

| **Area** | **x** | **y** | **z** | **k** | **T** |
|---|---|---|---|---|---|
| Supplementary motor area | -6 | 0 | 56 | 30263 | 15.76 |
| R Primary motor cortex | 36 | -16 | 60 | | 14.95 |
| L Primary motor cortex | -40 | -38 | 48 | | 18.18 |
| L Superior parietal lobule | -32 | -52 | 54 | | 14.57 |
| R Superior parietal lobule | 34 | -48 | 56 | 7306 | 11.47 |
| R Cerebellum superior | 28 | -62 | -26 | 10442 | 14.28 |
| R Cerebellum posterior | 28 | -68 | -52 | | 11.81 |
| L Cerebellum superior | -18 | -54 | -24 | | 14.11 |
| L Cerebellum posterior | -20 | -64 | -48 | | 13.35 |
| L Putamen | -24 | 2 | 4 | 5102 | 12.46 |
| R Putamen | 26 | 2 | 4 | | 11.56 |

*Significance level set at p<0.05 corrected on the whole brain using family-wise error correction (FWE). K represents cluster size (# of voxels).*

#### Comparisons between experimental groups, activations outside regions of interest

*Table S6: Functional imaging results for Session 2 SRTT practice: between-group comparisons (a,b) and PPI analyses (c-e) based on seed regions identified in (a,b).*

| Area | x | y | z | k | T |
|---|---|---|---|---|---|
| **a. Main effect of schema compatibility (COMP > INCOMP)** | | | | | |
| R Calcarine sulcus | 20 | -72 | 6 | 1366 | 4.07 |
| L Cuneus | -8 | -92 | 26 | 25 | 3.72 |
| L Medial occipital cortex | -28 | -76 | -2 | 20 | 3.45 |
| L Lingual gyrus | -14 | -50 | 0 | 31 | 3.41 |
| **b. Main effect of schema incompatibility (INCOMP > COMP)** | | | | | |
| L Anterior frontal cortex | -44 | 48 | 16 | 27 | 3.51 |
| **c. Psycho-physiological interaction, seed region left primary motor cortex (INCOMP > COMP)** | | | | | |
| R Frontal eye fields | 2 | 34 | 50 | 179 | 4.32 |
| R Medial temporal gyrus | 52 | -34 | -16 | 29 | 3.64 |
| R Superior frontal cortex | 30 | 66 | 8 | 76 | 3.73 |
| L Superior frontal cortex | -26 | 68 | 2 | 27 | 3.66 |
| L Fusiform gyrus | -64 | -54 | -4 | 31 | 3.54 |
| L Anterior prefrontal cortex | -40 | 56 | -8 | 8 | 3.44 |
| R Medial temporal gyrus | 54 | -6 | -22 | 7 | 3.36 |
| L Medial temporal gyrus | -62 | -20 | -12 | 9 | 3.34 |
| **d. Psycho-physiological interaction, seed region right primary motor cortex (INCOMP > COMP)** | | | | | |
| R Occipital cortex | 24 | -92 | 8 | 7 | 3.36 |

*Only activations outside regions of interest are reported here, refer to Table 3 in the main text for activations within regions of interest. Significance level set at $p<0.001$ uncorrected.*

### Neuroimaging results: multivariate pattern analyses

#### Exploratory analyses on declarative memory schema ROIs

We conducted exploratory multivoxel pattern similarity analyses of two ROIs traditionally associated with the schema memory model in the declarative domain, i.e. the angular gyrus (Wagner et al., 2015) and the medial prefrontal cortex (mPFC) (van Kesteren et al., 2013). The results of the *group* x *ordinal position* ANOVAs are reported in Table S7 and corresponding Figure S1. In both ROIs, there was a significant main effect of ordinal position, but no main effect of group or interaction of group and ordinal position. These results suggest that these regions are not sensitive to the schema-compatibility of the motor learning context. Thus, although both the mPFC and angular gyrus were shown to play a key role in schema-mediated learning in the declarative domain (Bonnici et al., 2012; Frankland & Bontempi, 2005; Takashima et al., 2006, 2007; Wagner et al., 2015), our results suggest that they are not involved in the corresponding process in the motor domain.

*Table S7: Results of exploratory multivariate pattern analyses on declarative memory schema ROIs for the 2 groups (COMP/INCOMP) x 8 ordinal positions (1 through 8) ANOVAs.*

| ROI | Main effect Group | | | | Main effect Ord | | | | Group x Ord interaction | | | |
|---|---|---|---|---|---|---|---|---|---|---|---|---|
| | df | F | $p_{corr}$ | Part $\eta^2$ | df | F | $p_{corr}$ | Part $\eta^2$ | df | F | $p_{corr}$ | Part $\eta^2$ |

| | | | | | | | | | | | | |
|---|---|---|---|---|---|---|---|---|---|---|---|---|
| Ang | 1,57 | 0.25 | 0.62 | 0.004 | 4.42,252.19 | 16.76 | **<0.001*** | 0.227 | 4.42,252.19 | 1.06 | 0.38 | 0.018 |
| mPFC | 1,57 | 0.00 | 0.97 | 0.000 | 4.02,229.06 | 3.11 | **0.02*** | 0.052 | 4.02,229.06 | 2.11 | 0.08 | 0.036 |

*Ang = angular gyrus; mPFC = medial prefrontal cortex; df = degrees of freedom; part η2 = partial eta squared. Significant p-values are marked with an asterisk and bold font.*

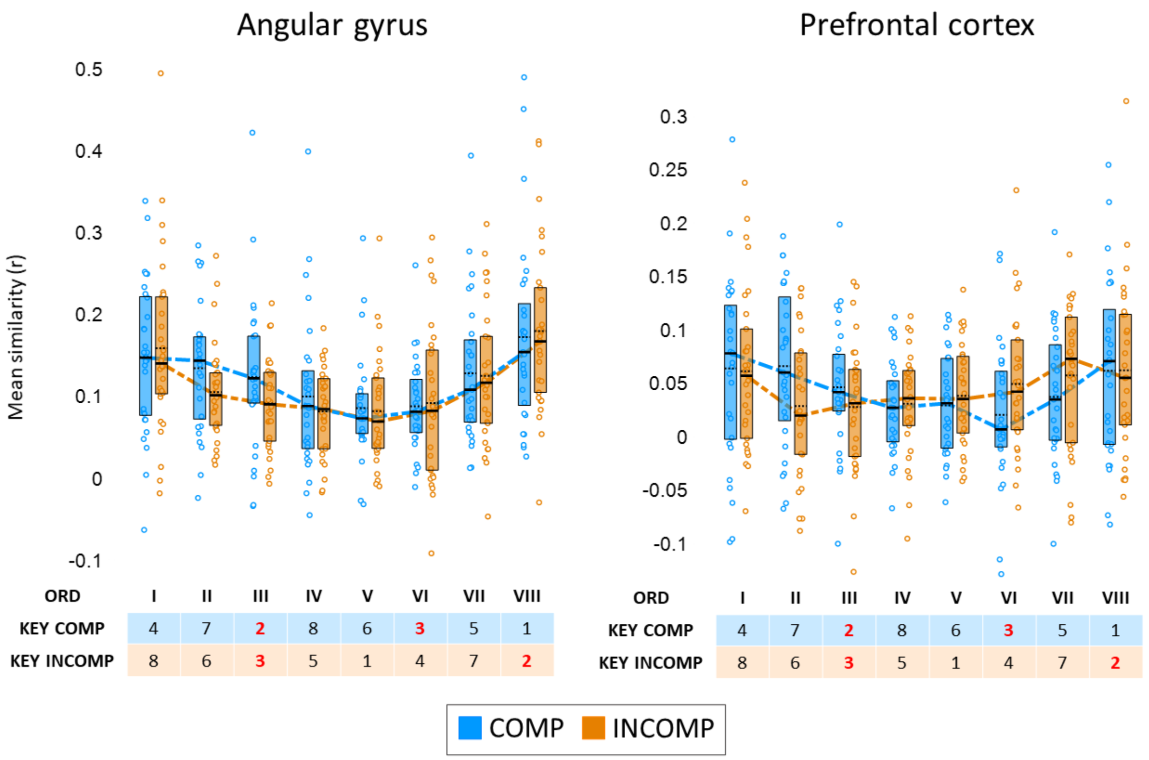


*Figure S1: Mean pattern similarity per key/ordinal position pairing, for each experimental group and exploratory ROI. Black horizontal bars (full) indicate the median and black horizontal bars (dotted) the mean. Boxes represent the interquartile range (IQR). Coloured dashed lines connect the medians in each experimental group. Coloured circles represent individual data.*

#### Supplementary material – additional references


Destrebecqz, A., Peigneux, P., Laureys, S., Degueldre, C., Fiore, G. Del, Aerts, J., Luxen, A., Van Der Linden, M., Cleeremans, A., & Maquet, P. (2005). The neural correlates of implicit and explicit sequence learning: Interacting networks revealed by the process dissociation procedure. Learning & Memory, 12(5), 480–490. https://doi.org/10.1101/LM.95605

Lehéricy, S., Bardinet, E., Tremblay, L., Van De Moortele, P. F., Pochon, J. B., Dormont, D., Kim, D. S., Yelnik, J., & Ugurbil, K. (2006). Motor control in basal ganglia circuits using fMRI and brain atlas approaches. Cerebral Cortex, 16(2), 149–161. https://doi.org/10.1093/CERCOR/BHI089

Penhune, V. B., & Doyon, J. (2002). Dynamic Cortical and Subcortical Networks in Learning and Delayed Recall of Timed Motor Sequences. Journal of Neuroscience, 22(4), 1397–1406. https://doi.org/10.1523/JNEUROSCI.22-04-01397.2002

Penhune, V. B., & Doyon, J. (2005). Cerebellum and M1 interaction during early learning of timed motor sequences. NeuroImage, 26(3), 801–812. https://doi.org/10.1016/J.NEUROIMAGE.2005.02.041

Strange, B. A., Fletcher, P. C., Henson, R. N. A., Friston, K. J., & Dolan, R. J. (1999). Segregating the functions of human hippocampus. Proceedings of the National Academy of Sciences, 96(7), 4034–4039. https://doi.org/10.1073/PNAS.96.7.4034
